# Vegetation clump size and number as indicators for alternative stable states in semi-arid ecosystems

**DOI:** 10.1101/2025.10.03.680294

**Authors:** Z. Chen, K. Halder, J. A. Bonachela

## Abstract

Semi-arid ecosystems are vulnerable to desertification, a pressing issue in the face of global climate change due to the expected increase in aridity in drylands. In these ecosystems, vegetation often grows in spatially regular patterns that differ as aridity increases (gaps, labyrinths and clumps), which has been widely documented empirically and studied theoretically aiming to understand how these patterns relate to desertification. While some theoretical models able to reproduce these regular patterns predict smooth desertification transitions, most typically predict an abrupt transition linked to the possibility of two alternative stable states (desert and vegetated states), predictions that are yet to be confirmed empirically. If this bistability of alternative stable states occurs, however, environmental fluctuations and the history of the ecosystem determine which state ultimately materializes, thus making it harder to predict desertification and, therefore compounding the challenges posed by it. Here, we combine empirical data and theoretical methods to investigate the links between bistability and vegetation spatial organization, aiming to identify indicators for the presence of alternative stable states. We found that the presence of vegetation clumps is not necessarily indicative of bistability, but how their morphology responds to aridity can provide reliable indicators of bistability and an impending desertification transition. Our results show that a decrease in the size or number of clumps does not necessarily precede desertification, and that monitoring clump size as aridity changes helps differentiate between ecosystems in which alternative stable states emerge from purely local vegetation-water interactions versus those in which they emerge from feedbacks occurring at different scales. By identifying bistability and its underlying mechanisms, our methodology indirectly identifies whether desertification will occur abruptly, whether restoration efforts should consider a potential ecosystem history-dependence, and offers potential targets for ecosystem management.

## Introduction

The survival of an ecosystem often depends on the delicate balance between biotic and abiotic factors. Climatic change affecting these factors can lead to large qualitative changes in the ecosystems, a phenomenon referred to as an ecological transition. Ecological transitions can be observed at any scale, and are widely (and increasingly) reported in the literature using terms such as critical transitions, catastrophic shifts, or tipping points (Lenton et al., 2023). The socio-economic and ecological reverberations of such transitions can be profound, as they can lead to decreased biodiversity and loss of key ecosystem functions. That is the case of desertification transitions, where increased aridity can shift a dryland ecosystem from a vegetated state to a barren one. Desertification hotspots are home to more than 500 million people as of 2015, and future climate is expected to expand drylands to up to three billion people (Intergovernmental Panel on Climate Change [IPCC], 2022; Rosenzweig et al., 2014). The increase in aridity will therefore exacerbate the scope and magnitude of desertification processes, threatening biodiversity and land-based assets, agricultural production and, more generally, the economic activity of the impacted regions (IPCC, 2022; Rey et al., 2017; Walther, 2010). Thus, given that drylands at risk of desertification are at a crucial nexus of ecosystem services, functioning, human development, and biodiversity concerns, identifying ways to assess ecosystem health and help describe and predict desertification is a pressing task of crucial importance.

In arid and semi-arid ecosystems, where the average precipitation only meets less than 50% of potential plant water demand, the temporal and spatial variation of vegetation is highly influenced by water availability (Snyder and Tartowski, 2006). In these environments, as aridity increases so does plant competition for water, which ultimately leads to desertification. Thus, water is the main growth-limiting resource and driver for desertification transitions. In addition to competition for water, occurring where root systems overlap, plant growth is facilitated locally as the root system breaks soil biocrust thus improving water infiltration, and plant shading contributes to preventing water evaporation. This short-range facilitation and long-range competition (mechanisms referred to as scale-dependent feedbacks, SDF; Rietkerk and van de Koppel (2008)) have been shown theoretically to generate regular vegetation patterns found in semi-arid ecosystems as aridity changes (Deblauwe et al., 2008; Borgogno et al., 2009). Thus, the spatio-temporal response of vegetation to changes in aridity has been studied theoretically to assess ecosystem health and identify potential indicators to predict desertification.

By invoking SDF, very different mathematical models have successfully reproduced the sequence of vegetation patterns observed in arid and semi-arid ecosystems as rainfall decreases (gaps, labyrinths, and clumps, see insets in Fig. 1) (Klausmeier, 1999; HilleRisLambers et al., 2001; von Hardenberg et al., 2001; Lejeune, 2002; Okayasu and Aizawa, 2001; Rietkerk et al., 2002; Gilad, 2004; Ursino, 2005, 2007; Siero, 2020; Martinez-Garcia et al., 2023). In these models, clumped vegetation patterns emerge for the driest condition, ultimately leading to desertification if rainfall keeps decreasing (Deblauwe et al., 2011), evidencing a link between vegetation self-organization and the desertification transition.

**Figure 1:**
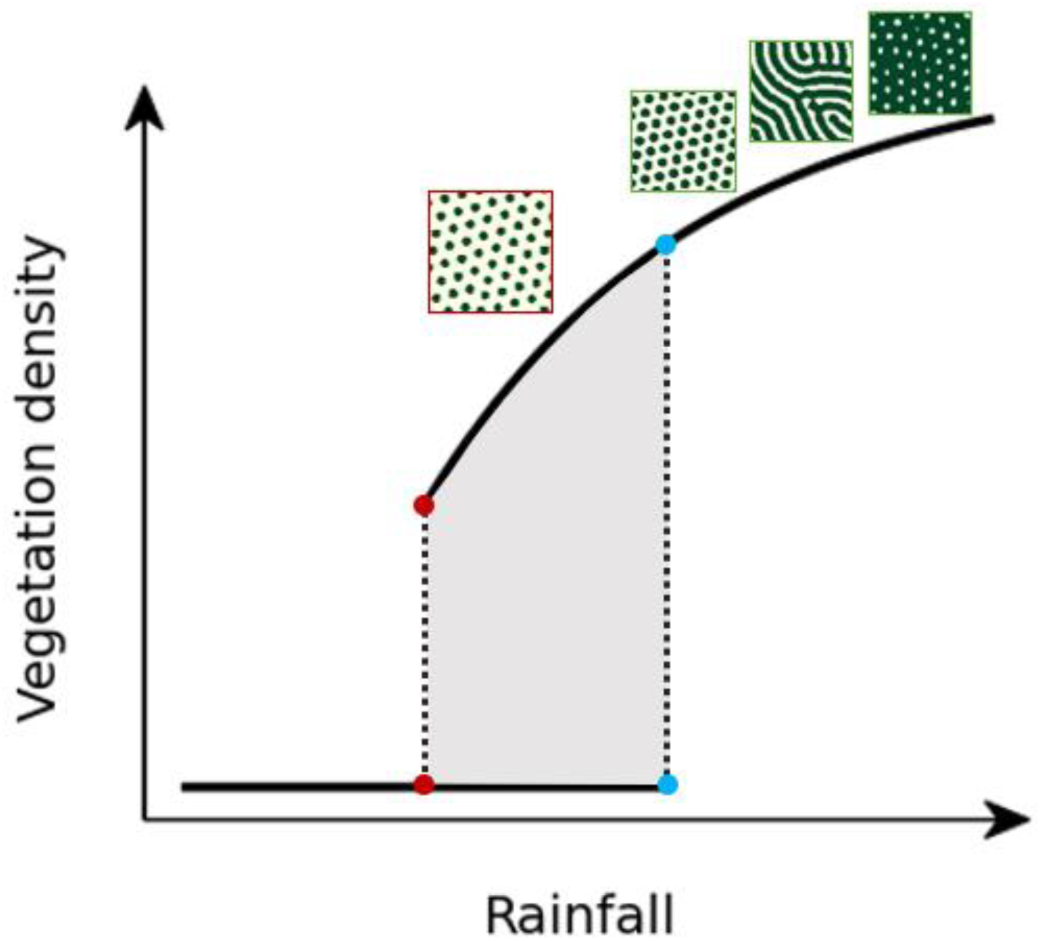
Schematic diagram illustrating a discontinuous vegetation transition in response to changes in rainfall. The transition features a bistability regime with simulated patterns: the blue dots mark the start of the bistability regime (shaded area), while the red dots indicate where the system ultimately transitions to the desert state. Because of bistability, however, the transition may occur in the rainfall range between the two dashed lines.

While some models predict a smooth, progressive transition, most models predict an abrupt desertification and thus a potentially catastrophic transition (Fig.1) (Martinez-Garcia et al., 2023). Moreover, recent mathematical one-dimensional approaches suggest that vegetation reorganizes spatially to increase resilience in a changing environment, which could ultimately lead to either smooth or abrupt transitions depending on whether the reorganization is successful (Siteur et al., 2014; Bastiaansen et al., 2018; Vanderveken et al., 2023); the generality of these results, however, has been recently contested (Pinto-Ramos & Martínez-García, 2026). Importantly, the lack of data has prevented empirical corroboration of any of the predictions above. Determining the shape of the transition is nonetheless important because, theoretically, discontinuous transitions are accompanied by bistability and hysteresis, which would make desertification even more difficult to predict and revert. Hysteresis (tendency for the system to follow different paths depending on its history i.e., its previous states) means that the rainfall levels needed to restore the system after desertification (blue points in Fig.1) would be higher than the rainfall levels for which the system became a desert (red points, same figure). In addition, bistability means that the system can potentially reach either one of two alternative stable states (or even switch dynamically from one to the other) for the same value of the environmental variables: in Figure 1, the system could be either vegetated or desertic within the rainfall range between the red and blue points, with the realized state decided by the initial states or environmental fluctuations (Lejeune et al., 2002; Lejeune et al., 2004; Meron et al., 2004; Sherratt and Lord 2007; Kéfi et al., 2010). Therefore, within the wide bistability range that precedes desertification, rainfall levels do not unequivocally predict ecosystem state (e.g. desertification can occur even if rainfall is far from the red dots in Fig.1, the so-called desertification tipping-point) and restoring past environmental conditions does not necessarily ensure ecosystem regeneration. Thus, given the uncertainty and ecological consequences associated with bistability, determining whether an ecosystem is within a bistability regime is crucial for environmental management and conservation.

Existing empirical data and theoretical studies have traditionally focused on developing methods to identify and monitor ecological transitions. To our knowledge, however, no existing method can account for or utilize spatial self-organization to determine whether the system is within a bistability regime. The sequence of patterns as rainfall decreases, and clumps more specifically, have generally been proposed as early-warning signals for the desertification transition (Rietkerk et al., 2004; Scheffer et al., 2009; Dakos et al., 2011). However, clumps can be found within and out of the bistability range (Fig. 1). Moreover, the analysis of pattern morphology using Fourier transforms (Deblauwe et al., 2011) has been used in theoretical studies to anticipate desertification: the distance between clumps (and therefore the wavelength where the fundamental peak of the Fourier transform occurs, see Supplementary Information) has been shown to increase as the transition is approached (Siteur et al., 2014). However, whether such wavelength changes can be related to bistability is unknown. Enhance vegetation index (EVI) time-series and other remote-sensing products have been used to measure temporal and spatial indicators such as correlation, skewness, and variance that behave in characteristic ways as the transition approaches (Veldhuis et al., 2022). Nonetheless, these indicators (e.g., critical slowing down) have not been linked to bistability, require local perturbation analyses that are labor-intensive in the field, and/or require long time series that capture the transition (Dakos et al., 2011).

Importantly, the technical and conceptual challenges above have also prevented confirmation of an aspect as fundamental as to whether the desertification transition is abrupt (i.e., catastrophic) or smooth. This information, and the associated ecological consequences (e.g., the potential presence of hysteresis), can be substantially helpful for managers and decision-makers (Deblauwe et al., 2011; Martinez-Garcia et al., 2023). The presence of bistability could corroborate indirectly the presence of an impending catastrophic transition.

Here, we aim to fill this gap by studying the relationship between vegetation patterns and bistability. We analyzed pattern morphology from satellite images using a study region in Sudan, and a model tailored to that region, to better understand the differences across resulting vegetation patterns and how those differences relate to bistability. We parameterized the model using available data for our focal system, calculating key parameters such as plant and water diffusivity, typically estimated qualitatively in the literature. We used Fourier transforms to monitor pattern regularity, and found a previously overlooked feature — a secondary fundamental peak — that can capture the changes in main properties of the vegetation spatial distribution as well as anticipate the transition. We link the changes in this peak as rainfall decreases to changes in clumps that are unique to the bistability regime, therefore pointing to simple metrics that can be used to identify the bistability regime (if it exists) and thus assess ecological uncertainty in the system. We tested our predictions using satellite images from our focal region, and observed similar pattern behavior to those observed within the bistability regime of the model. Thus, according to our predictions the focal area would be undergoing bistability and shows behavior indicative of an impending abrupt desertification transition. Our methodology, potentially general to any system with SDFs, can help assess the risk of transition and ecosystem health for ecosystems where spatial self-organization occurs, and therefore can potentially be used to design and test management policies for such ecosystems.

## Methods

### Study area

Our study area is located in the western part of South Kordofan in Sudan (Fig. 2; 11° 42’ 0’’-11° 0’ 0’’ N, 27° 54’ 0’’-28° 0’ 0’’E; (Esri, 2009)), part of a larger region where vegetation patterns have been reported in the past (Deblauwe et al., 2011). Topographically, the area has an overall slope of 0° (ArcGIS Online at a 25 m spatial resolution (Esri, 2013)). At the effective plant root zone depth, the soil texture is loam and the root zone plant-available water holding capacity ranges between 55 and 83 mm (Africa SoilGrids online resources at a 1 km spatial resolution (Leenaars and Hengl, 2015; Leenaars and Hengl, 2017)). Thus, there is no significant difference across the study area in terms of slope and soil properties (Supplementary Figure S1.1).

**Figure 2:**
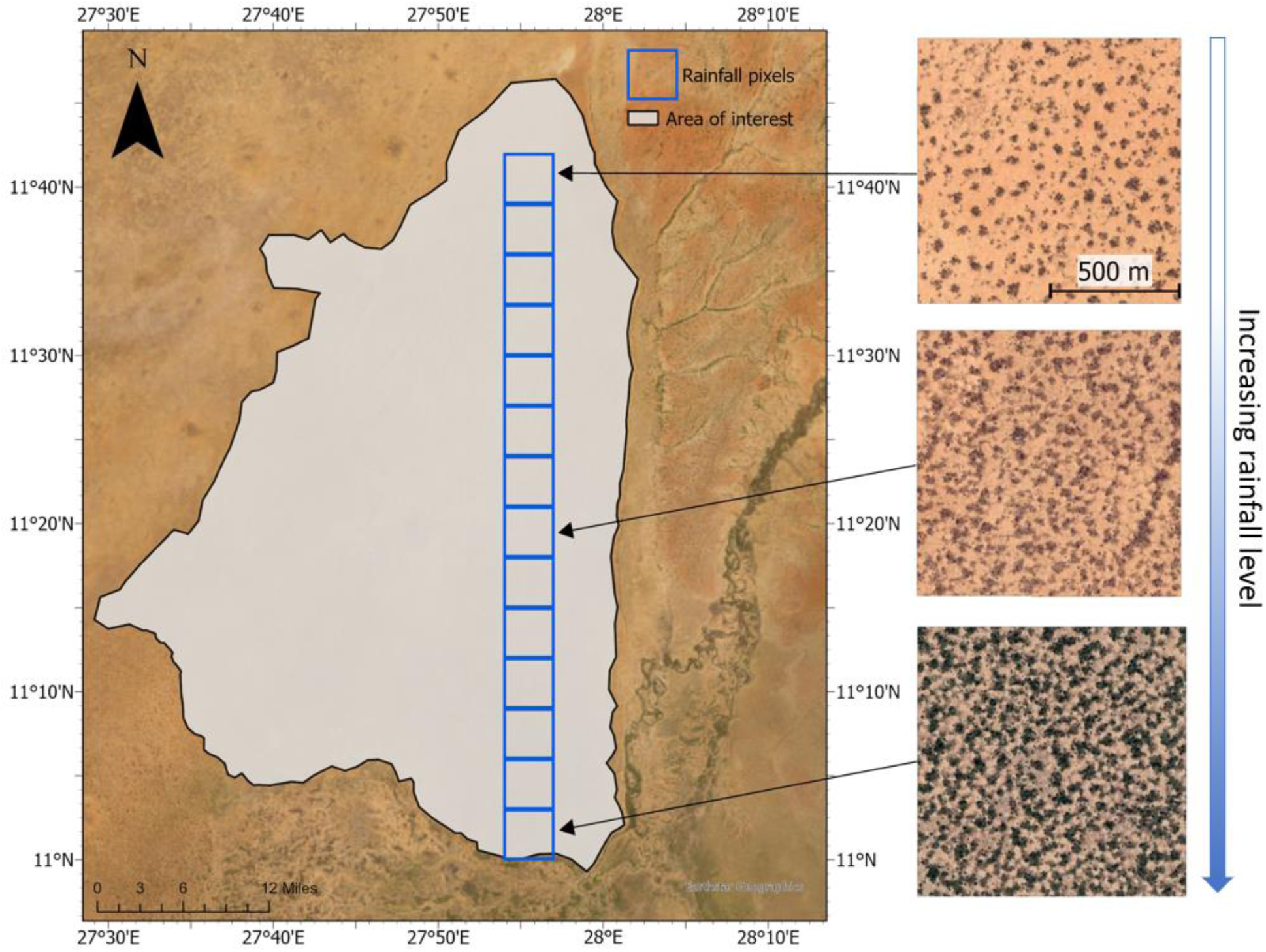
Optical satellite image of study area and different patterns across rainfall levels. The blue squares delineate the rainfall pixels from CHIRPS Precipitation Data (1990-2020). The selected 1km×1km vegetation patterns on the right are illustrative examples of clumps, labyrinths, and labyrinthine-gap-like (from top to bottom).

The mean annual precipitation in our study area ranged between 220 and 590 mm over the last three decades (CHIRPS Precipitation Data: 1990-2020), with a monthly rainfall level that decreases from south to north. As rainfall decreases, vegetation patterns change from gap-like to labyrinths to clumps (Fig. 2). We selected 14 rainfall pixels on the same longitude (27° 54’ 0’’-28° 0’ 0”E), which ranged these different types of patterns.

The information above justified focusing on rainfall as the main environmental feature driving the observed changes in patterns.

#### Focal vegetation species

We focused on *Acacia mellifera*, a native shrub in semi-arid regions in western Asia and Africa including Sudan (CABI, 2022). Various *A. mellifera* patterns have been reported in our particular focal area Kordofan, Sudan (Wickens and Collier, 1971; Deblauwe et al., 2011). *A. mellifera* tends to release its seeds in a small radius around the parent shrub and this aggregated dispersal can result in clump-like aggregated growth consisting of different age groups (Hagos, 2001).

### Modified Klausmeier model

To represent our system, we chose the generalized version of one of the simplest and earliest models for self-organized vegetation patterns in semi-arid ecosystems, the Klausmeier model (Klausmeier, 1999; Ursino, 2005). The model describes the dynamics of water density (*w*; *kg_H_*_2*O*_*m*^−2^) and plant density (*p*; *kg_drymass_m*^−2^):

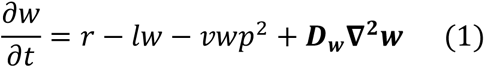

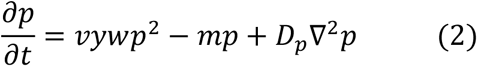

The first term in Eq. (1) represents precipitation (only water input in this model), which occurs at rate *r* (*kg_H_*_2*O*_*m*^−2^*year*^−1^). The second and third terms represent, respectively, water loss due to evaporation and transpiration and plant growth water uptake, where *l* (*year*^−1^) is the water loss rate and *v* ((*kg_drymass_m*^−2^)^−2^*year*^−1^) is related to water maximum uptake rate. The last term represents water spatial movement, here encoded as diffusion at a rate *D_w_* (*m*^2^*year*^−1^). This diffusive term differentiates this from the original Klausmeier model, which instead considered directional water movement (e.g., due to slope terrain, represented with an advection term, that is, a first spatial derivative). The first term in Eq. (2) represents plant biomass growth, where *y* represents the yield of plant biomass per unit water consumed, in *kg_drymass_*(*kg_H_*_2*O*_)^−1^. The second term represents plant natural mortality (at a rate *m*; *year*^−1^), the only cause of plant biomass loss considered here. The last term in the equation represents plant dispersal, which occurs at a rate *D_p_* (*m*^2^*year*^−1^). The model considers that water uptake and growth benefit from the presence of other plants nearby (increased infiltration and reduced evaporation), reason why those terms are quadratic in *p*.

Without dispersal of water and plants, i.e. in the non-spatial, mean-field case, the model results in homogeneous (i.e. non-patterned) systems, with desert being the only possible outcome (i.e. only stable state) for rainfall below 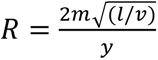, and bistability of vegetation and desert for larger rainfall values (Klausmeier, 1999). With dispersal, i.e. when spatially explicit interactions are considered, the model has been reported to generate all three types of regular vegetation patterns (Ursino 2005). Therefore, this model has the potential to replicate the typical patterns observed in semi-arid regions with a reduced number of parameters. Thus, to ensure that any pattern resulting from the model is representative of our study area, we next found tailored values for these parameters and, in particular, for the dispersal (i.e. diffusivity) parameters.

### Model parameterization

Values for the parameters of the model were obtained directly from the literature or through calculations tailored to our focal region.

#### Water-related parameterization

To replicate the gradient of rainfall within the study area, we explored fixed values of the rainfall parameter *r* between a realistic range based on the monthly rainfall data obtained from CHIRPS Precipitation Data. We alternatively implemented a stochastic, dynamic rainfall function generated from the monthly rainfall data, aiming to mimic the seasonality within the focal semi-arid climate (see SI Section 2 for details).

The water loss rate (*l*) was obtained by dividing the theoretical water evapotranspiration for one year by the yearly water availability (precipitation). To calculate the theoretical water evapotranspiration, we used the Thornthwaite Potential Evapotranspiration equation (see SI Section 3a, Eq. S3.2) for Al Nahud and Muglad, two cities with data collection that overlapped with the study area (Thornthwaite, 1948; Alam et al., 2013). The estimated water loss rate range was from 3.7 to 5.4 *year*^−1^ (see SI Section 3a for detailed calculation).

On the other hand, models for vegetation in arid/semi-arid ecosystems often follow a qualitative approach to parameterizing water diffusivity (*D_w_*) or reuse values from classic work (Sherratt, 2016). Here, we calculated water diffusivity aiming to keep the same thoroughness as with the rest of ecological parameters. To this end, and under the assumption that *w* represents surface water, we calculated the estimated range of water diffusivity by using the classic Manning equation for the velocity of a liquid (Manning, 1891), considering flow velocity in all directions. As a result, we estimated a maximum theoretical water diffusivity and a more conservative value within the same order of magnitude, *D_w_* = 170 *m*^2^*year*^−1^, which is also in line with the few estimates available in the literature. See SI Section 3a for details.

#### Vegetation parameterization

For the yield of plant biomass per unit water consumed (*y*), we used the standard assumption in the literature for encroacher species (*Terminalia sericea* and *A. mellifera*) of 1500 liters of water consumed to produce 1 kg of dry mass (NAU, 2010). For the mortality rate (*m*), the existing literature provided a range of values for *A. mellifera* with 0.03 and 0.54 *year*^−1^ as extremes (see SI section 3b).

On the other hand, we interpreted diffusivity parameters for plants (*D_p_*) as the successful dispersal event for the focal species while considering germination rate and first-year seedling survival rate. Our calculations provided a plant diffusivity between 1.15-1.88 m^2^ per year (see SI Section 3b for detailed calculation). To acknowledge that plant dispersal also results from shrub growth (i.e. clonal reproduction), which happens more slowly, we used a more conservative value within that order of magnitude (*D_p_* = 1 *m*^2^*year*^−1^).

Finally, the parameter accompanying the water uptake term (*v*) had units that are not typical for an uptake rate ((*kg_drymass_m*^−2^)^−2^*year*^−1^), and therefore we could not find a value following ecological arguments as above. Instead, we used a value within the range in the original model, which was calibrated for semi-arid ecosystems (Klausmeier, 1999). Our choice (*v = 57*) resulted in patterns within rainfall ranges comparable to those observed in the focal area. Given that this is the only parameter that we could not constrain using ecological arguments, we also ran a sensitivity analysis, confirming that other values of the parameter did not qualitatively alter our conclusions. See section S3 for further details and Supplementary Table S3.2 for final parameterization.

### Simulations

We ran our simulations in a 250×250 spatial mesh, using periodic boundary conditions to reduce finite-size and border effects. The spatial resolution (*dx*) was 4 meters (i.e., simulated area of 1 km×1 km), and the time resolution (*dt*) was 0.001 year. The value of *dx* was small enough to resolve pattern motifs with several pixels, and large enough to cover a big area that provided a statistically adequate number of those motifs. We used the Jacobi method to discretize the two-dimensional Laplacian that represents diffusion, and the Euler method to integrate the partial differential equations of our model due to its simplicity and ease of implementation (especially under a stochastic environment, see above).

Following the standard protocol, we started simulations at the homogeneous stable state for the given rainfall value; we further perturbed randomly chosen locations representing a small percentage of the system by adding a certain amount of plant biomass (Table S2). We then let the system run a minimum of 100,000 years to allow stable patterns to emerge. Note that this maximum time depends on simulation details, i.e. is not necessarily characteristic of the real system. We ran a minimum of 25 replicates and stored final snapshots of the plant density matrix for further analysis.

### Pattern analysis

#### Image preparation

We analyzed the patterns observed in snapshots from our simulations, as well as 1km×1km optical satellite images with 1.2 m spatial resolution for the focal study area extracted from Google Earth Pro. Because we lacked information about biomass density for the focal area, to ensure a meaningful comparison with the model we first converted all the images to black-and-white scale, which is a standard step in the analysis of vegetation patterns. Thus, we analyzed images only based on the presence or absence of vegetation at a given pixel. For this binarization, we used a standard unsupervised classification algorithm (Kmeans, see details in SI section 4).

#### Power spectrum analysis

To study changes in spatial regularity, we calculated the radially averaged power spectrum associated with each snapshot collected at each rainfall level. We first scaled the binarized images to ensure zero mean and unit variance across pixel intensities and examples, then obtained the two-dimensional (discrete) Fast Fourier Transform (2D FFT) using the well-established Python algorithm in the NumPy package (Harris et al., 2020). We further computed the radially averaged power spectrum by averaging across all radii from the center of the 2D power spectrum in all directions, normalized by their respective bin-width areas. Peaks in this radially averaged power spectrum denote regularity in the original pattern (see SI section 5).

#### Clump size and count distributions

As justified in results, measuring clump size and number can be informative when the rainfall levels are close to catastrophic levels. In order to measure the size and number of clumps from snapshots, we used the two-pass Hoshen–Kopelman algorithm from the SciPy Python library to identify and label each clump. For the binarized versions of both the simulation and satellite images, we calculated the average number, size, and maximum size over replicates for each rainfall level.

Finally, adjoining clumps that are still in the process of separating and forming independent ones can occur for rainfall levels for which patterns are formed more slowly, even at very long simulation times. To avoid any bias introduced by these temporary motifs, we isolated and excluded them using an unsupervised Isolation Forest algorithm (Liu et al., 2008). See SI section 4 for further details.

### Other vegetation models

We also assessed which features observed with the tailored model (if any) are specific to bistability, that is, to the simultaneous coexistence and/or stability of both desertic and vegetated states for the same environmental conditions. Here, we distinguish between bistability observed when space is not accounted for (i.e. for the equations without the diffusion terms, hereon *mean-field bistability*) and bistability observed in the spatially explicit system (i.e. equations with water and vegetation diffusivity, hereon *spatial bistability*). To this end, we qualitatively compared the results we obtained with the Klausmeier model (which shows both mean-field and spatial bistability) to those obtained with another model for regular vegetation patterns that does not show any bistability and instead predicts a smooth desertification transition (Martinez-Garcia et al., 2013). In this comparison, we also included a model with only spatial bistability (Rietkerk et al., 2002), and a version of that model that shows both types of bistability but in a more limited range than the Klausmeier model (Siero, 2020). See SI section 6 for further details.

#### Further analysis of bistability

For each model, we determined the range of rainfall levels for which mean-field bistability and spatial bistability (if present) occur.

For the determination of mean-field bistability, we used classic analytical methods to determine the ranges of rainfall for which a space-less version of the equations representing the dynamics of the system show stability for both the desertic and the vegetated state. For the focal model, for example, this method involved analyzing Eqs. (1)-(2) without the spatial derivatives looking for its stationary states, then calculate their corresponding eigenvalues to check whether there are values of rainfall for which both stationary states show stability (all eigenvalues are negative); as explained above, bistability of homogeneous desertic and vegetated states occurs for rainfall levels above 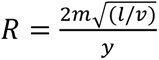.

To determine for which ranges patterns can emerge in the full, spatial system, we followed two approaches. First, we found the rainfall values for which initializing the system with a high or a low addition of plant density to random locations (10% of the pixels) leads to different final states, i.e. patterns versus desert, respectively. This method provided the realized spatial bistability region, which is dependent on details of the simulation such as how the initial condition is perturbed. Second, we analytically assessed the stability of each periodic solution to disturbances that slowly alter the spacing of the pattern itself, i.e. the so-called Eckhaus mechanism (Eckhaus, 1965). The region defined by the Eckhaus instability contours is also known as the Busse balloon (Siteur et al., 2014). This method thus identifies the maximum theoretical range of bistability for the model, defined as the rainfall levels between the lower Eckhaus instability contour and the beginning of mean-field bistability. See SI section 5 for summary of the method, reported in detail in (Halder and Bonachela, 2026a).

## Results

In our study area, different levels of annual rainfall led to different types of patterns (Fig. 3a-3c). The clump pattern occurred in areas with rainfall levels of 430.8 to 463.1 mm/year, while the labyrinth pattern was observed in areas with rainfall levels of 463.6 to 481.4 mm/year. We found dense labyrinths that also showed features of the gap pattern in areas with rainfall levels ranging from 496.0 to 508.9 mm/year. We measured the average percentage of each snapshot that is covered by vegetation, as a proxy for vegetation density. The representation of this metric against rainfall levels can be used to indicate whether the system is in a vegetated or desertic state, thus providing the so-called phase diagram of the system (see schematic example in Fig.1). For our study area, however, the phase diagram did not clearly indicate the character of the expected desertification transition nor whether there is bistability (Fig. S1.2), which justified the use of a model tailored to our focal system to provide a more complete picture of the transition.

**Figure 3:**
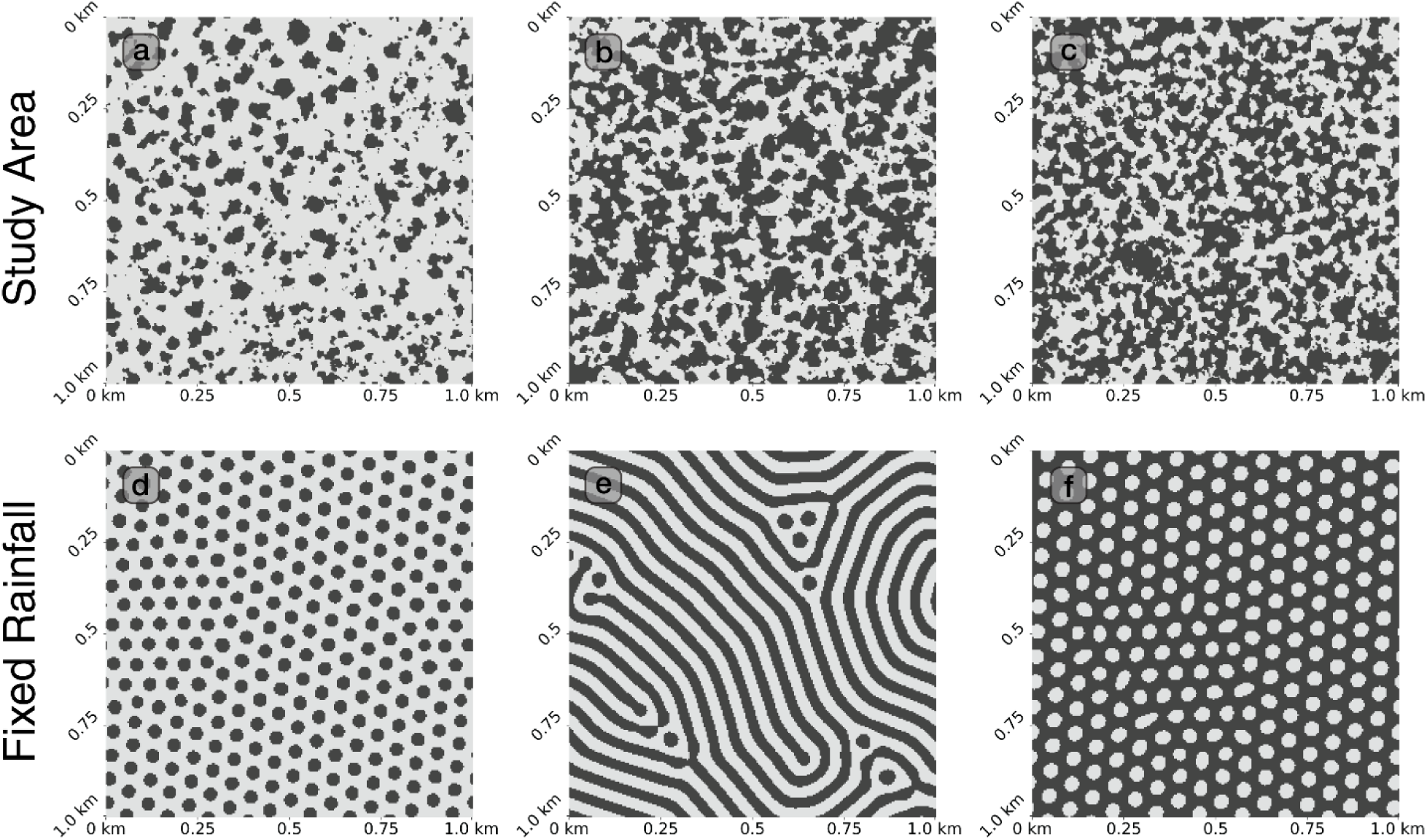
Binarized versions of the real vegetation patterns from our study area and simulated patterns. Panels (a), (b) and (c) are satellite images derived from Google Earth pro after binarization (i.e., black-and-white transformation, see Methods), with annual precipitation 430.8, 478.0 and 508.9 mm/year, respectively. Panels (d), (e) and (f) correspond to the simulation with annual precipitation *r* = 430, 475 and 510 mm/year, respectively. For all snapshots, black indicates the presence of our focal vegetation and white means bare soil.

### Simulated vegetation patterns and phase diagram

With the parameterization described in Methods and exploring different (fixed) precipitations, our simulations of the Klausmeier model were able to generate clump, labyrinth, and gap patterns at similar rainfall levels to those observed in the study system (Fig. 3d-3f). The patterns obtained with a data-informed stochastic rainfall function (see SI section 2 and Fig. S2.1) were very similar to those obtained with a fixed rainfall level (Fig. S2.2). Thus, for simplicity we used fixed rainfall levels for our subsequent simulations and analyses.

We first studied the desertification transition by constructing the phase diagram. To that end, we first calculated the stationary values of vegetation and water for the mean-field system (i.e. without considering water or vegetation dispersal). As the solid lines in Fig. 4 show, mean-field bistability of the desertic and vegetated states occurred for rainfall levels above *R* = 444 mm/yr. Next, we obtained the phase diagram for the complete, spatially explicit model by measuring the stationary value of the vegetation density (spatial average of the density across pixels for a given snapshot, averaged across replicates) for a given rainfall level (points in Fig. 4). Given the sensitivity to initial conditions expected for systems with bistability, to obtain the spatially explicit stationary states we used two different initial perturbation values (1.0 *kgm*^−2^ and 0.1 *kgm*^−2^). Both perturbations resulted in similar patterns for rainfall levels under 375 and over 442 mm/year: the desert state was observed for rainfall levels under 375 mm/year; clumps, when rainfall ranged from 443 to 460 mm/year; labyrinths were observed when rainfall ranged from 460 to 510 mm/year; and gaps were found when rainfall ranged from 510 to 525 mm/year. However, the two perturbations led to very different results from 375 to 442 mm/year: the simulation with an initial perturbation value of 1.0 *kgm*^−2^ generated clump patterns, while the simulation with an initial perturbation value of 0.1 *kgm*^−2^ resulted in extinction. The spatially explicit system with our tailored parameterization, thus, showed spatial bistability between 375 and 442 mm/year (spatial-bistability regime).

**Figure 4:**
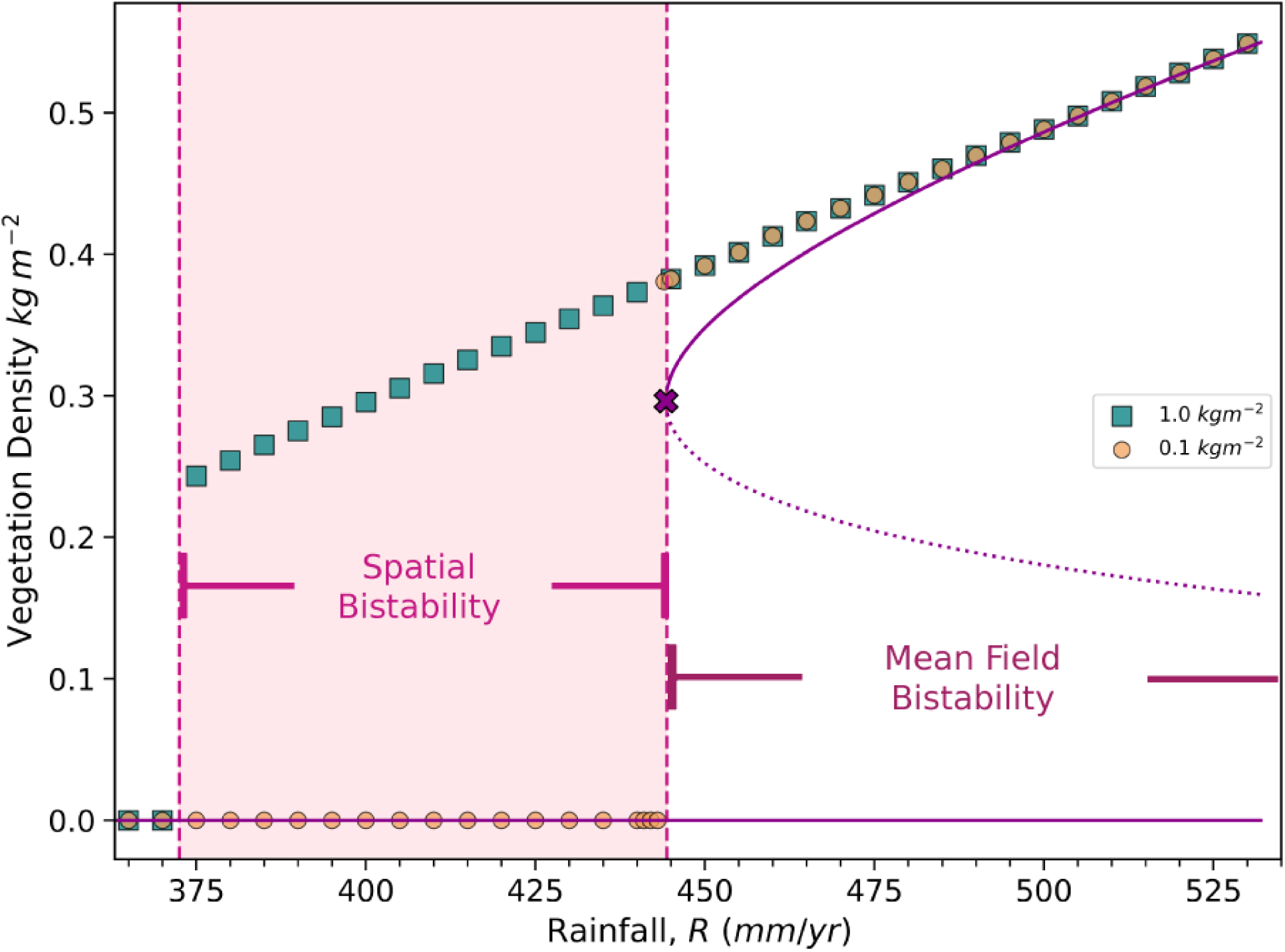
Phase diagram obtained by representing the values of mean vegetation density in the spatially explicit model (symbols) and the stationary states for vegetation in the mean-field (i.e. non-spatial) version of the model (lines) as a function of rainfall levels. The spatial bistability region starts at 370 mm/yr, whereas the mean-field bistability region starts at 444 mm/yr.

### Radially averaged power spectrum

To understand if the properties of the patterns (e.g., regularity) changed across the rainfall gradient, we studied how the power spectrum for each simulated replica’s final snapshot changed with rainfall. As expected, the heatmap of the power spectrum obtained across rainfall levels after averaging over replicates showed a primary fundamental peak for all patterns, with spatial frequencies ranging between 15 and 19 cycles/km (red dashed line, Fig. 5). The primary fundamental peak represents the wavelength of the motif that is repeated in the pattern. For the clump pattern, for example, the increase of spatial frequency with increasing rainfall availability signifies shorter distances between nearest neighboring clumps. Also as expected, the power spectrum exhibited harmonic peaks at twice and thrice the spatial frequency of the primary fundamental peak for all simulated rainfall levels. As explained in Supplementary Information section 5.2, for spatial-bistability rainfall values, the location of this primary fundamental peak falls within a region of the space of wavenumbers where theory predicts the emergence and stability of spatial patterns even if the non-spatial (mean-field) model would predict only the stability of the desertic state (Busse balloon region, defined between *R* = 340 mm/yr and *R* = 530 mm/yr, see Fig. S5.6). The maximum range of spatial bistability would then be defined by rainfall between 340 mm/yr and 444 mm/yr; importantly, the wavenumber of the observed patterns closely follows the upper boundary of this region.

**Figure 5:**
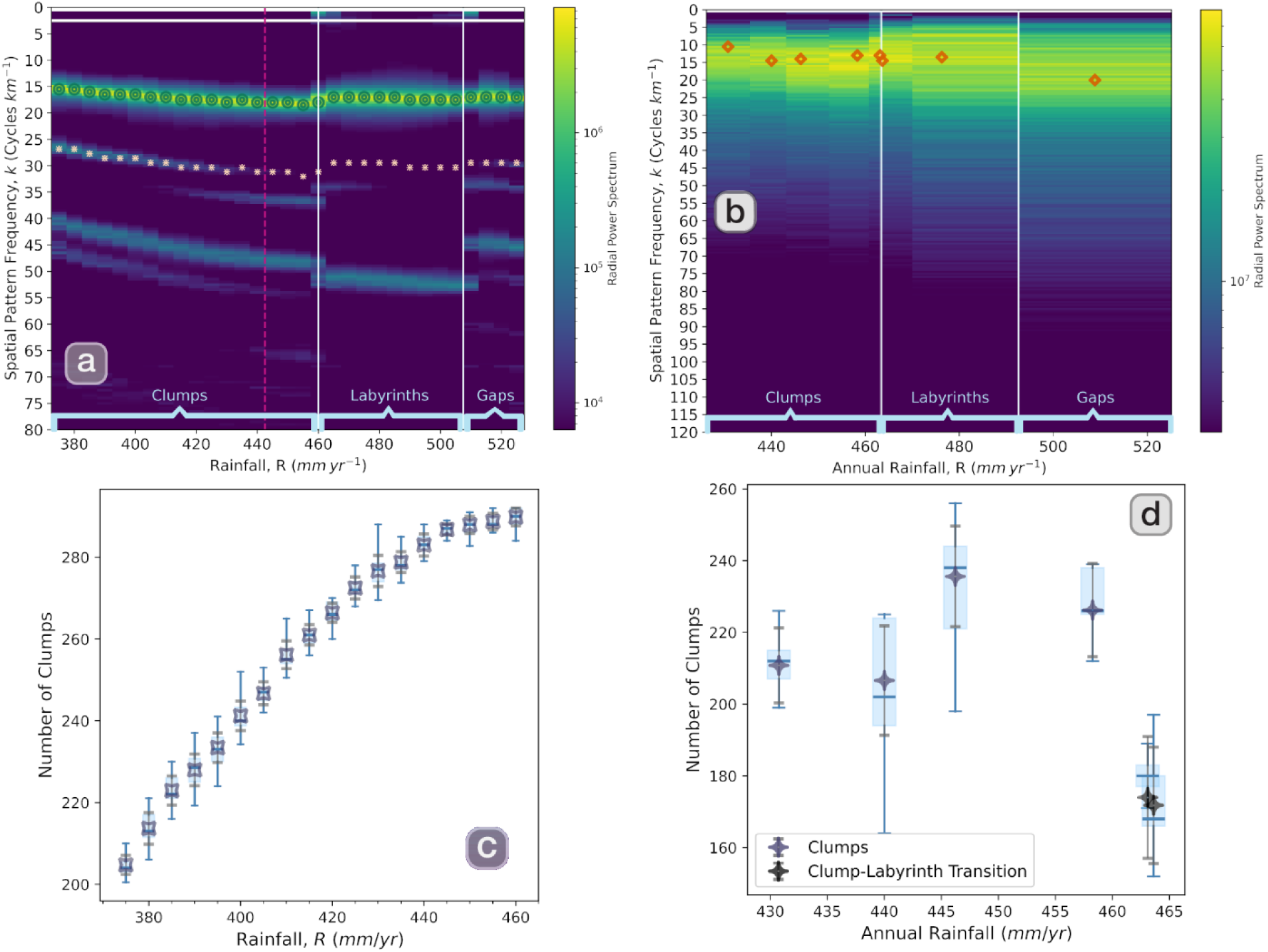
Upper row: heatmaps of the power spectra for patterns obtained from our simulations (panel a) and study site (panel b). Coordinates are the spatial frequency of the pattern and rainfall levels. Color represents the value of the radially averaged power spectrum for the given spatial frequency (see definition in SI; yellow means highest and dark blue lowest). For simulations, the first and second fundamental peaks are highlighted with green dots and pink asterisks, respectively; for the study site, only the first fundamental peak was noticeable (red diamonds). Light blue lines were used to separate different types of patterns. For simulations, the vertical dashed line separates the spatial bistability regime (375-442 mm/year). Bottom row: average number of clumps measured in snapshots of the simulation (panel c) and study site (panel d) across replicates. Symbols represent the mean value, with grey error bars denoting ±1 standard deviation across replicates; boxes span the interquartile range, with whiskers extending to 1.5× the interquartile range and horizontal line in box representing median.

Surprisingly, the semi-log plot also revealed a second fundamental peak located around 26 cycles/km (pink asterisks, Fig. 5). This secondary peak occurred when the underlying pattern was clumps or gaps, but not labyrinths. As explained in section 5, this peak is related to the distance between next-nearest neighbors of a given motif. Due to the hexagonal symmetry of clump and gap patterns, the peak is located at a wavenumber that results from multiplying by 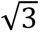 that of the primary fundamental peak, and can be seen as a mixed mode of the Fast Fourier Transform. Inside the bistability regime, the height of this secondary fundamental peak decreased as rainfall increased (see colors in heatmap), growth that was accompanied by an increase in height of the first harmonic of the fundamental peak. See SI section 5 for details.

The relationship between this changing secondary peak and the spatial bistability regime make it a potential indicator for bistability and even an impending desertification. However, the heterogeneity and stochasticity associated with real vegetation patterns and environmental conditions prevented this secondary fundamental peak or any of the harmonics from being discernible in the power spectra of images from our study area (Fig. S5.7). Thus, to understand better the phenomenology above and whether it was specific of the Klausmeier model or a general signature of bistability, we repeated the analysis with the models in (Martínez-García et al., 2013; Rietkerk et al., 2002; Siero, 2020), all of which produce similar vegetation patterns but make different predictions regarding the desertification transition with and without considering space (Figs. S6.1, S6.4, S6.7). As shown in Figs. S6.2, S6.5, S6.8 and summarized in table 1, for all models the radial spectrum showed a secondary fundamental peak that followed a similar qualitative behavior as rainfall decreases. However, in the model without spatial bistability (Martinez-Garcia et al., 2013), the peak was noticeable for all the clump-pattern region, and the location of the primary fundamental peak (and therefore the distance between clumps in the pattern) did not change with growth conditions in the model with a smooth desertification transition.

**Table 1:** Summary of results obtained from analyzing the focal study area (South Kordofan), the main model tailored to this focal area (tailored Klausmeier model), and three other models that show no bistability (Martinez-Garcia et al model), only spatial bistability (Rietkerk model), or both types of bistability (Rietkerk model with evaporation).

|  | **Behavior as aridity increases<br>(i.e. as rainfall decreases)** |  |  |  |  |
|---|---|---|---|---|---|
|  | **Spatial bistability** | **Mean-field bistability** | **Average clump size** | **Number of clumps (alternatively, primary FFT peak)** | **Secondary FFT peak** |
| **South Kordofan (real system)** | – | – | Constant | Declining | – |
| **Tailored Klausmeier model** | Yes | Yes | Constant | Declining | Height increasing from negligible |
| **Rietkerk model** | Yes | No | Declining | Declining | Height increasing from negligible |
| **Rietkerk model with evaporation** | Yes | Yes | Declining then constant | Declining | Height increasing from negligible |
| **Martinez-Garcia model** | No | No | Declining | Constant | Height increasing from non-negligible value |

### Clump size and number change with rainfall

Although the noise typically observed for power spectra calculated from remote-sensing images makes it difficult (if not impossible) to all the radial spectrum phenomenology described above, as justified in SI sections 5 and 6 such phenomenology is closely related to changes in clump size and number. Therefore, the behavior of pattern clump size and count across examples indicates a potential more direct pathway to identify bistability.

We measured and compared clump size and count across rainfall levels for our simulations and study area. In all cases, and as predicted by symmetry arguments in SI section 5, the number of clumps was proportional to the wavelength where the primary fundamental peak is located. For the tailored Klausmeier model (Fig. 6a-b), outside the spatial bistability region the average clump size decreased as rainfall decreased, whereas the number of clumps remained constant. In the spatial bistability region, however, the net change in clump size and maximum clump size remained small, whereas the number of clumps decreased with rainfall. We observed a similar behavior for the other model that shows both spatial and mean-field bistability (Siero, 2020) (Fig. S6.9). The model with only spatial bistability (i.e. no mean-field bistability) (Rietkerk et al., 2002) showed decreasing clump size and number as rainfall decreased (Fig. S6.6). Finally, for the model with a smooth transition (Martinez-Garcia et al 2013), clump size decreased with declining growth conditions whereas the number of clumps remained constant (Fig. S6.3). These observations therefore indicate differences in clump pattern behavior experienced in the presence and absence of spatial and mean-field bistability.

**Figure 6:**
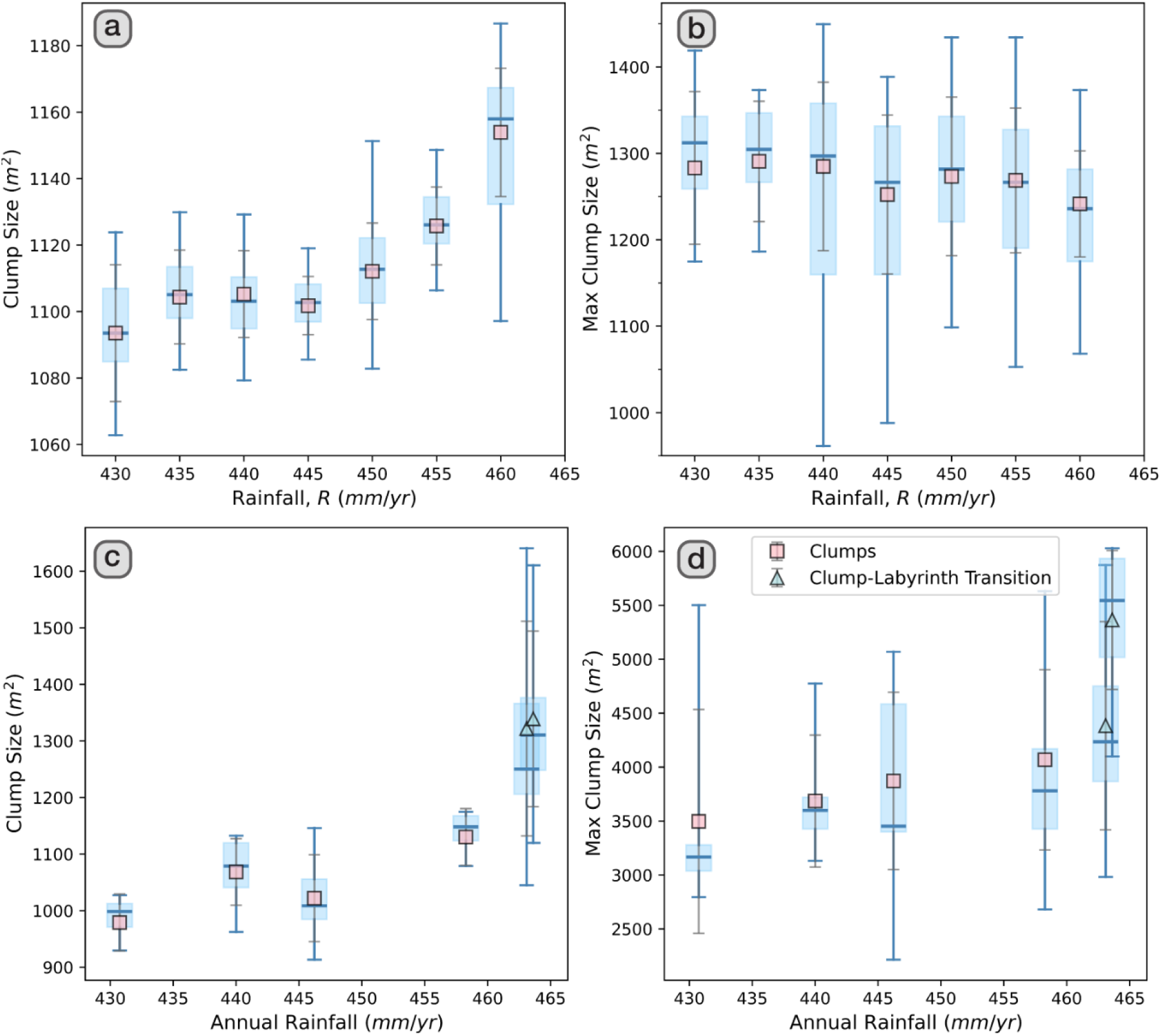
Comparison of average clump size (left column) and maximum clump size (right column) as a function of rainfall levels for simulations (upper row) and study site (bottom row). Note that “clump-labyrinth transition” corresponds to patterns that converge (very slowly) to labyrinths but still showed some clumps when the simulation was stopped. For easier comparison, we focused the range of rainfalls in simulations to the ones observed in the study site; see SI section 6 for results from a variety of other models.

For satellite images (Fig. 6c-d, and S6.3a-c), clump size slightly increased as rainfall increased, then jumped to higher values after clumps transitioned into labyrinths. The number of clumps increased with rainfall levels but decreased for labyrinthine patterns. As expected, although the behavior of the number of clumps was somewhat approximated by the wavelength of the primary fundamental peak in the radial spectrum, the approximation was not as close as in any of the models above. Importantly, except for the highest rainfall level snapshot, the location of these peaks fell within the Busse balloon region defined by our tailored version of the modified Klausmeier model (diamonds in Fig. S5.6).

## Discussion

Semi-arid ecosystems are home to hundreds of species, for which desertification can have catastrophic consequences. Thus, the increased risk of desertification associated with expected future climatic changes threatens food and economic security of the millions of people who inhabit these ecosystems. Aiming to understand better how desertification occurs, here we combined remote-sensing data and a classic theoretical model to explore whether it is possible to identify signatures of bistability in the spatial response of vegetation as aridity increases.

### A tailored, ecologically grounded parameterization for the modified Klausmeier model

We used here a modified version of the original Klausmeier model that replaces a directional advection term for water movement with a diffusion term. The latter was needed as the data we compiled for our study area indicated a region with little to no slope, indicating no directional water movement.

Existing work has utilized this modification to the Klausmeier model, or similar models, to study vegetation patterns, typically in one spatial dimension. Here, the comparison with remote-sensing images of our study area required a focus on two dimensions. The reduced number of parameters in this model allowed us to inform it with a bottom-up parameterization approach by which we obtained tailored, ecologically relevant parameter values, then studied whether patterns emerged for realistic rainfall levels. We identified a native plant species (*A. mellifera*) that is highly likely to underlie the observed vegetation patterns, and derived ranges for the parameters in Eq. (S2)-(S4) that corresponded to this species. For plant diffusivity, for example, we considered successful plant dispersal events by taking into account seed dispersal rate, outside canopy ratio, germination rate, clonal reproduction, and seedling survival rate. Regarding water diffusion, there is surprisingly very little information in the vegetation pattern modeling literature about the physical processes involved, and thus very little justification for the values typically used for water diffusivity (Sherratt, 2016). The water variable in the Klausmeier model, for example, does not discern between superficial or subterranean water. This distinction is important because, ultimately, only underground water can be utilized by the root system. Here, we chose to interpret the water variable as superficial water and assume perfect water infiltration (i.e., all superficial water is available for the root system), which facilitated the calculation of the diffusion coefficient. Thus, we drew inspiration from the Manning equation for the velocity of a liquid and calculated a plausible range of diffusion rates. Despite the importance of the diffusivity parameters for vegetation pattern models, there is no definition or standardized way to calculate them, with further difficulties stemming from a lack of clarity regarding the definition of plant or water diffusion in these models. The calculations we used for our parameterization provide an ecological justification and a range that can be used to constrain parameter values in future theoretical work. Nonetheless, interdisciplinary efforts in which theoreticians and empiricists agree on definitions for the different components of these models have the potential to provide clear instructions to measure these important parameters (Martinez-Garcia et al., 2023).

Our parameterization for the model resulted in emerging patterns that were comparable to those of the study area, which served as an initial qualitative validation.

### Clumps and bistability can be characterized through previously unidentified features of the power spectrum

The radially averaged power spectrum is typically used to differentiate between types of patterns based on the existence or location of a primary fundamental peak (Deblauwe et al., 2011). This was also true in our study, as the location of the primary fundamental peak jumped whenever patterns changed from clumps to labyrinths or from labyrinths to gaps (more subtly for real-world images). In addition, we further identified a second fundamental (i.e., non-harmonic) peak. Regardless of whether this is a real fundamental peak or a mixed Fourier mode of the pattern, this peak and the primary fundamental peak showed a distinctive behavior when the system is bistable: an increase in rainfall caused an increase in the location of the primary fundamental peak, and the decrease in height (and eventual disappearance) of the second fundamental peak. As justified in SI, the latter can be linked to changes exclusive to the spatial bistability region; the only model with a smooth desertification transition showed instead a constant primary peak and a declining but non-negligible secondary peak as growth conditions improved.

These results together reveal that, when bistability is present, clumps converge to labyrinths in a distinct way that can be used to identify bistability. On the one hand, the different behavior of the primary fundamental peak observed in these models, identifies it as an indicator for bistability. Although the changes in this peak with rainfall have been suggested as an indicator of ecosystem health worsening as aridity increases (Rietkerk et al. 2004; Deblauwe et al. 2011), our analyses rule out this possibility since ecosystem deterioration (i.e. decreasing rainfall) does not necessarily translate into changes in the location of the primary fundamental peak for all the models simulated here. From a technical point of view, whether the location of the primary fundamental peak changes with rainfall depends on the existence of a region of the phase diagram where stable vegetation patterns emerge in the desertic region thanks to the spatial interactions present in the system (i.e. existence of spatial bistability and the Busse balloon region; Halder and Bonachela, 2026a), regardless of the presence of mean-field bistability. Moreover, the shape of the maximum spatial bistability region (i.e. the shape of the Busse balloon) conditions the behavior that the location of the primary fundamental peak follows within that region as rainfall changes, in this case leading to a decreasing frequency (increasing wavenumber/location of the primary peak) as rainfall decreased. On the other hand, our results reveal that the emergence and increase in height of the secondary fundamental peak is conserved across models and, therefore, this secondary peak can be used as an alternative early-warning indicator of impending desertification. Moreover, whether the secondary peak becomes negligible as rainfall increases flags the presence of spatial bistability. For real-world examples, however, the expected noise in the power spectrum but also the eventual disappearance of the peak limit the useability of this secondary peak: according to the heatmaps obtained for the tailored Klausmeier model, the secondary peak becomes negligible for *R* > 430 mm/yr, which is the smallest rainfall levels measured for our analyzed real-world images.

Importantly, the link between power spectrum and pattern features provides an understanding of the mechanisms underlying the behavior of the peak that enables bypassing the power spectrum altogether, and shifts the focus to properties that are easily measurable with satellite images.

### Clump size and count behavior are a signature of bistability and early-warning signals for desertification

As changes in clump number and size are the primary factors contributing to the changes in the fundamental power-spectrum peaks, and these are general properties that can be easily captured from snapshots of vegetation patterns, we examined these properties for both the simulations and satellite images.

Our analysis of simulated patterns with the tailored Klausmeier model shows that a decrease in rainfall leads to a reduction in clump spatial frequency (i.e., increase of wavelength) associated with a decrease in the number of clumps. Existing work has reported increasing pattern wavelength with aridity for a larger region containing our focal area in Sudan (Deblauwe et al., 2011), and occurring in the modified Klausmeier model as the system approached the desertification transition (Siteur et al., 2014; Bastiaansen et al., 2020). These observations refer to the location of the primary fundamental peak of the power spectrum, which we also observed moves to lower frequencies (i.e., higher wavelengths) as rainfall decreases, indicating more distance between clumps and therefore fewer clumps within the observation window. As explained in those references, as aridity increases fewer clumps can survive, with a decline in number of clumps that becomes more subtle if rainfall changes are slow and gradual and/or for less-regular patterns (like the ones expected in real-world scenarios). Moreover, we reported a negligible net change in average clump size and maximum clump size within the spatial bistability region. To the best of our knowledge, our study is the first to show a correlation between all those changes and the existence of a changeable second fundamental peak.

Changes in height for this second fundamental peak (and the first harmonic of the primary fundamental peak) seem to be a more noticeable reflection of the changes in clump size and number than the changes in location of the primary fundamental peak. Importantly, the behavior of these peaks and how it reflects on clump size and number can be used to identify both spatial bistability and mean-field bistability. Our results reveal that, across all models simulated here, if both types of bistability occur then clump size becomes constant as rainfall decreases within the spatial bistability region (i.e. rainfall range in which both the clump and desertic state are stable); outside of this region, clump size increases with rainfall levels. If spatial bistability occurs but mean-field bistability does not, clump size and number increase with rainfall within the spatial bistability region. Finally, if no bistability occurs, clump size increases with rainfall while the number of clumps remains constant. In other words, an increasing number of clumps with rainfall indicates the presence of spatial bistability, whereas a constant clump size close to the transition indicates also the presence of an underlying mean-field bistability and the proximity of desertification.

### Bistability relates to ecosystem response

These differences have important ecological implications and reveal very different ecosystem responses to climatic change. If a system shows mean-field bistability, local ecological interactions (in this case, local competition and facilitation) suffice to maintain the coexistence of the vegetated state within the desertic state. When only spatial bistability occurs, the fact that these interactions show scale separation (long-range competition vs short-range facilitation) becomes relevant to maintain such coexistence. As a result, the Klausmeier model and those in Siero (2020) and Rietkerk et al. (2002) describe systems that respond to an increase in aridity (i.e. decrease in rainfall) by reducing vegetation biomass per clump and losing clumps all together. We hypothesize that this behavior stems from an increased range of competition occurring as rainfall decreases, which in turn leads to a sudden loss of resilience or stability of the vegetated state at desertification. For the Klausmeier and the model in Siero (2020), however, local interactions are able to maintain coexistence, thus hindering the increase in long-range competition and leading to a constant clump size close to desertification. In a system described by the model with no bistability, competition range is fixed and thus worsening growth conditions barely affect clump numbers. Additional empirical work, however, is needed to test this hypothesis. Nonetheless, we are currently investigating the mechanisms underlying the differences between vegetation patterns in the presence or absence of bistability, which will be reported elsewhere.

### Evidence of spatial and mean-field bistability in our study site

The behavior of average and maximum clump size as well as that of the number of clumps can therefore potentially be used to assess ecosystem health in semi-arid systems. After applying the method above to the satellite images from our focal area, we observed a behavior resembling that obtained with the tailored Klausmeier model. The primary fundamental peaks measured across rainfall levels fell within the Busse balloon predicted with the Klausmeier model; clump counts showed a hump-shaped behavior with rainfall, whereas both maximum and average clump size showed a seemingly constant behavior. All together would indicate that the system is at a state similar to that of the tailored Klausmeier model mid-way into the spatial bistability region, when the secondary fundamental peak has already become negligible. Thus, with caution due to the apparent outlier observed in the real-system number of clumps at low rainfall, the conclusion would be that the study area would be within a spatial bistability regime. Consequently, the system would be at a heightened risk of desertification, as environmental changes and stochasticity may trigger the transition (i.e. a jump from the vegetated state to a desertic state) even if aridity levels are not unusually low or extreme. Indeed, the tailored Klausmeier model predicts desertification at substantially lower rainfall levels, and thus these indicators would warn of an impending desertification (and, consequently, call for management action) when there is still a chance to avoid transition or, at least, ameliorate its consequences.

### Identifying (or not) bistability helps the management of desertification transitions

The phenomenology observed here thus provides guidance to manage systems at risk of desertification by, for example, monitoring clump number and size over time (*in situ* or through satellite images) as rainfall levels change. If, as expected from climate change, rainfall decreases over the years, a constant number of smaller and smaller clumps would reveal that desertification is approaching, but the transition will be gradual, reversible and, therefore, manageable. A decreasing number of clumps would indicate spatial bistability, and therefore the possibility for environmental fluctuations to drive the system to desertification abruptly, with the transition being imminent if fewer clumps do not correlate with smaller clump sizes. In these cases, past history of the system becomes important because of the hysteresis associated with bistability, and management efforts should focus on controlling large external disturbances like grazing or human activities, which can tilt the system towards desertification (Kéfi et al., 2007; Halder and Bonachela, 2026b). Thus, our study provides a workflow that combines empirical and theoretical methods to investigate pattern morphology and its relationship with bistability and the health of the ecosystem. Importantly, because this workflow identifies whether the system is within a bistability regime (and, moreover, the type of bistability), it provides information about the character of the transition, underlying mechanisms, and path to recovery. Therefore, the information provided by our approach can be used to develop policies that encourage sustainable land use practices and protect the livelihoods of local communities from land degradation, or to adapt agricultural practices to better suit the changing conditions of the land.

### Limitations

Certain limitations of this study also call for caution when using our methodology, thus suggesting further research directions. Such limitations stem from missing information regarding the study area, as well as technical limitations.

First, the lack of high-resolution historical satellite images (including for our focal area) made temporal analysis impossible due to the lack of such images before the 1990s. Also, despite our efforts to ensure that all site replicates had similar environmental conditions (other than rainfall), other factors for which we did not have information such as grazing and soil nutrients could also influence the pattern. Additionally, we assumed that the main plant species underlying the patterns was the typically dominant and pattern-forming *A. mellifera,* but other species are present in the ecosystem, and calculated ranges for plant and water parameters based on proxy observations and data. Finally, we made important simplifying assumptions about both water and vegetation transport. All these aspects could be improved with ground-truthing observations and dedicated experiments *in situ*.

Our analysis of images carries its own limitations. We performed classification on visible-band composites (only imagery available to us at the required resolution), and as a consequence vegetation indices exploiting the near-infrared could not be used. Moreover, shrub shadow may be classified as canopy, which is likely to inflate apparent clump areas somewhat. Nonetheless, this potential artifact would act consistently across images, and therefore should not affect the trends reported here and used for our conclusions.

Regarding the model, the phenomenological simplified depiction of the water-vegetation interactions, although mathematically convenient, leads to ecological conundrums. For example, the uptake term depends on a parameter *v* that does not have ecologically meaningful units and, although to some extent can be interpreted as an uptake rate, this lack of meaning prevented us from using bottom-up arguments to tailor its value to our specific system. Importantly, our main results were not altered qualitatively when testing other values for this parameter, which only moved the desertification and pattern transitions along the rainfall axis. Other, more detailed models used to represent vegetation in semi-arid ecosystems, like the ones tested here to identify fingerprints of bistability, are arguably more realistic but also more difficult to parameterize due to the additional terms that they incorporate. One important aspect to keep in mind when using such models is that, although they may show similar vegetation patterns, the dynamical properties of the model can differ significantly (see above and SI section 6).

Finally, a common feature of vegetation models with bistability is that the rainfall level at which patterns first emerge in the upper branch of the phase diagram (red point in Fig.1) is determined by the associated perturbation strength (here, 1.0 kg m^-2^, see Methods). Our focal region did not cover a range of rainfall levels that included desertification, and thus we lacked information to parameterize this perturbation. Nonetheless, this parameter only affects the width of the bistability region, not the patterns or pattern transitions. Future empirical work is needed to directly link the pattern and associated power spectrum changes reported here to the actual desertification transition.

## Acknowledgements

We would like to thank R. Martínez-García for his comment on an early version of the manuscript and the code to run the model in Martínez-García et al. (2013). This work was supported by NSF grant DMS-2052616 to J.A.B.

## Supplementary Information

### Section 1. Study area information

Our focal area is located in the western part of South Kordofan in Sudan, shown in the World Topographic Map (Figure S1.1a; 11° 42’ 0’’-11° 0’ 0’’ N, 27° 54’ 0’’-28° 0’ 0’’E; Esri, 2013a). This area is part of a larger region where vegetation patterns have been reported in the past (Deblauwe et al., 2011). For the topographic information of our study area, the slope map (Fig. S1.1b) was derived from ArcGIS Online at a 25 m spatial resolution (Esri, 2013b). We derived soil water holding capacity (Fig. S1.1c) and soil texture data (Fig. S1.1d) at a 1 km spatial resolution from the Africa SoilGrids online resources (Leenaars and Hengl, 2015; Leenaars and Hengl, 2017).

The vegetation cover percentage was calculated for each rainfall pixel that showed clump-like patterns, which corresponded to rainfall levels ranging from 430 to 465 mm/year. Within each rainfall pixel, we took five replicates at a 1 km spatial resolution and calculated the mean vegetation cover percentage across the replicates. (Fig. S1.2). This phase diagram neither indicated nor ruled out the presence of bistability between vegetated and desertic states for any rainfall levels (i.e. did not fully resemble the phase diagram in Fig. 1 nor showed a smooth transition to zero vegetation density).

**Figure S1.1:**
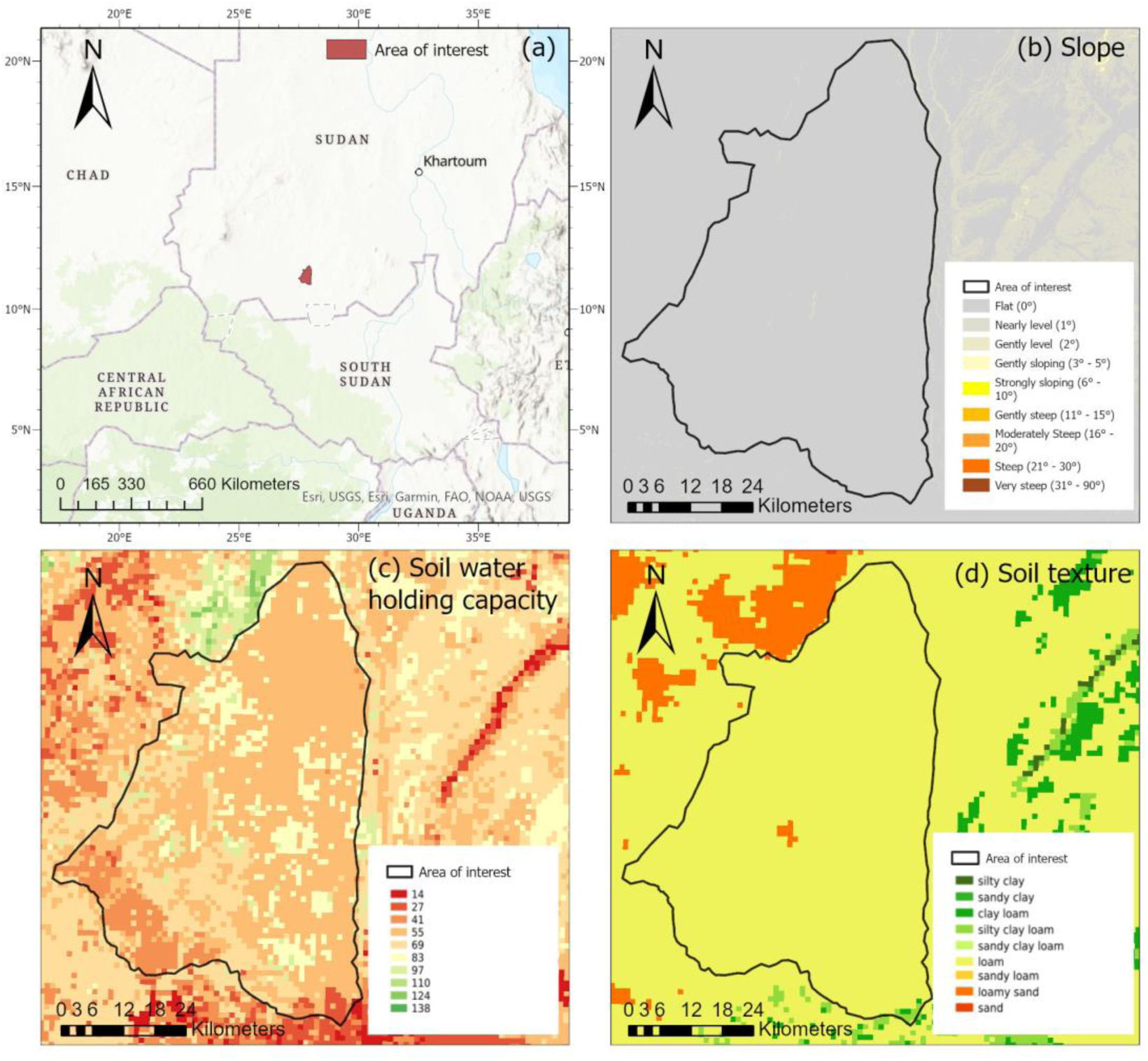
Focal area and main topographic properties. The study area (shown in panel (a)) is flat (slope between 0° and 1°, panel (b)). Panels (c) and (d) show that soil water-holding capacity and soil texture are similar at the effective plant-root zone depth across the region.

**Figure S1.2:**
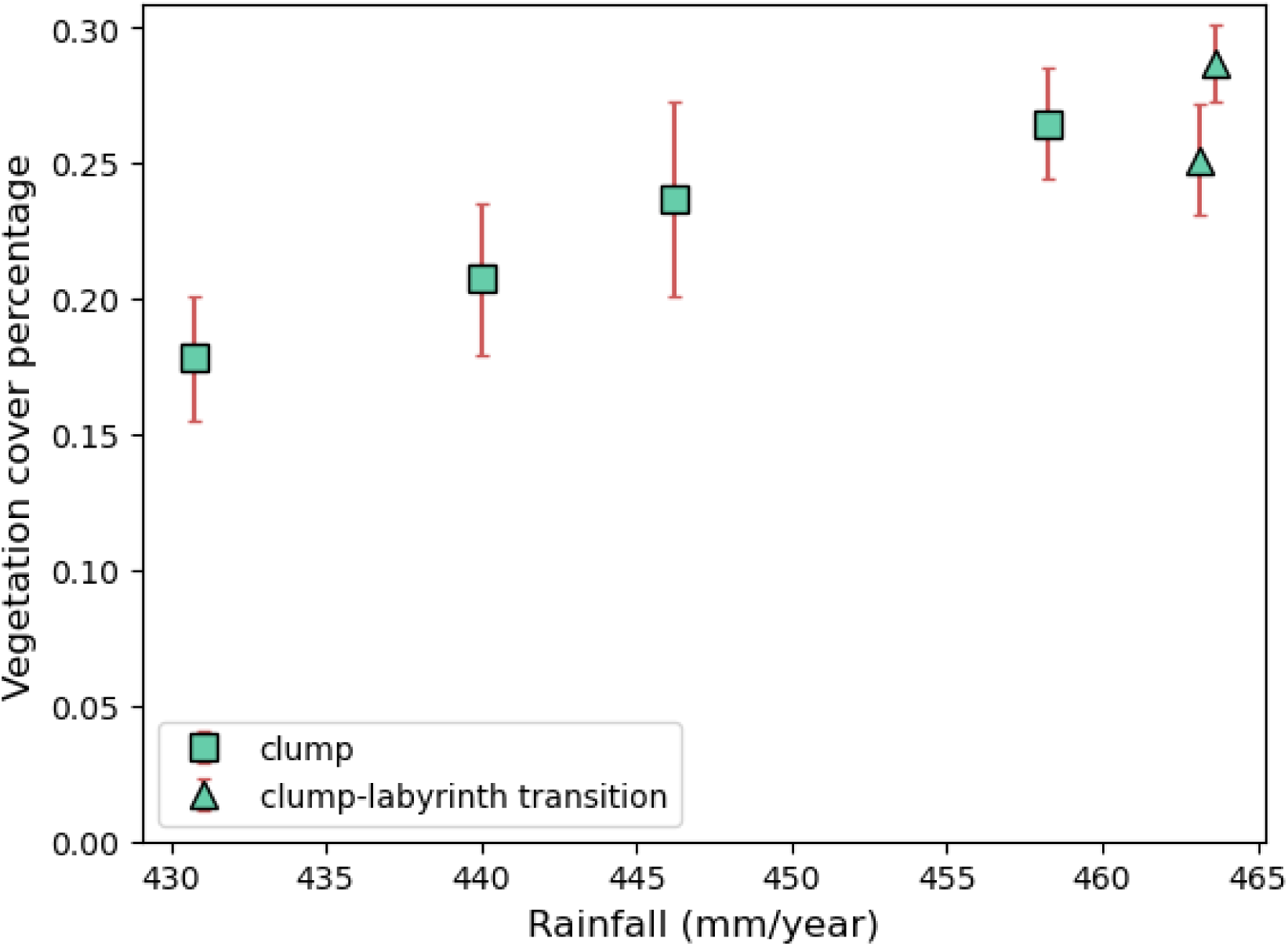
Phase diagram obtained representing the values of the mean vegetation cover percentage as a function of rainfall levels for our focal area. Error bars represent standard deviation with respect to the average across five replicates. Patterns that show both properties of clumps and labyrinth motifs are labeled using triangles.

### Section 2. Rainfall functions

To check whether simulated patterns are robust under stochastic and seasonal changes in rainfall, we replaced the constant value used for the rainfall parameter with a stochastic rainfall function generated from the monthly rainfall data. This function allowed us to resolve seasonality within the focal semi-arid climate, as well as the inherent climatic fluctuations.

To implement the seasonality part of the function, we first classified pixels according to the observed pattern (clump, labyrinth, or gap). We then estimated the medians and interquartile ranges of monthly precipitation over a 30-year period, which we fitted to a skewed-Gaussian function that provided us with a seasonal resolution of rainfall. The resulting stochastic rainfall function was (see Fig. S2.1):

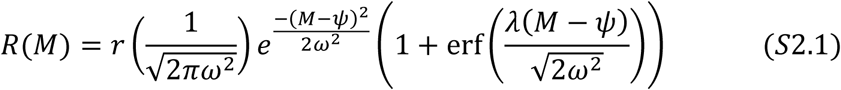

Where *r* is annual precipitation rate (mm/year), *M* is the specific month of the year, *ω* = 1.8, *ψ* = 8.8 and *λ* = −1.5.
To represent the stochasticity inherent to rainfall, we added a noise component to Eq. (S2.1).

**Figure S2.1:**
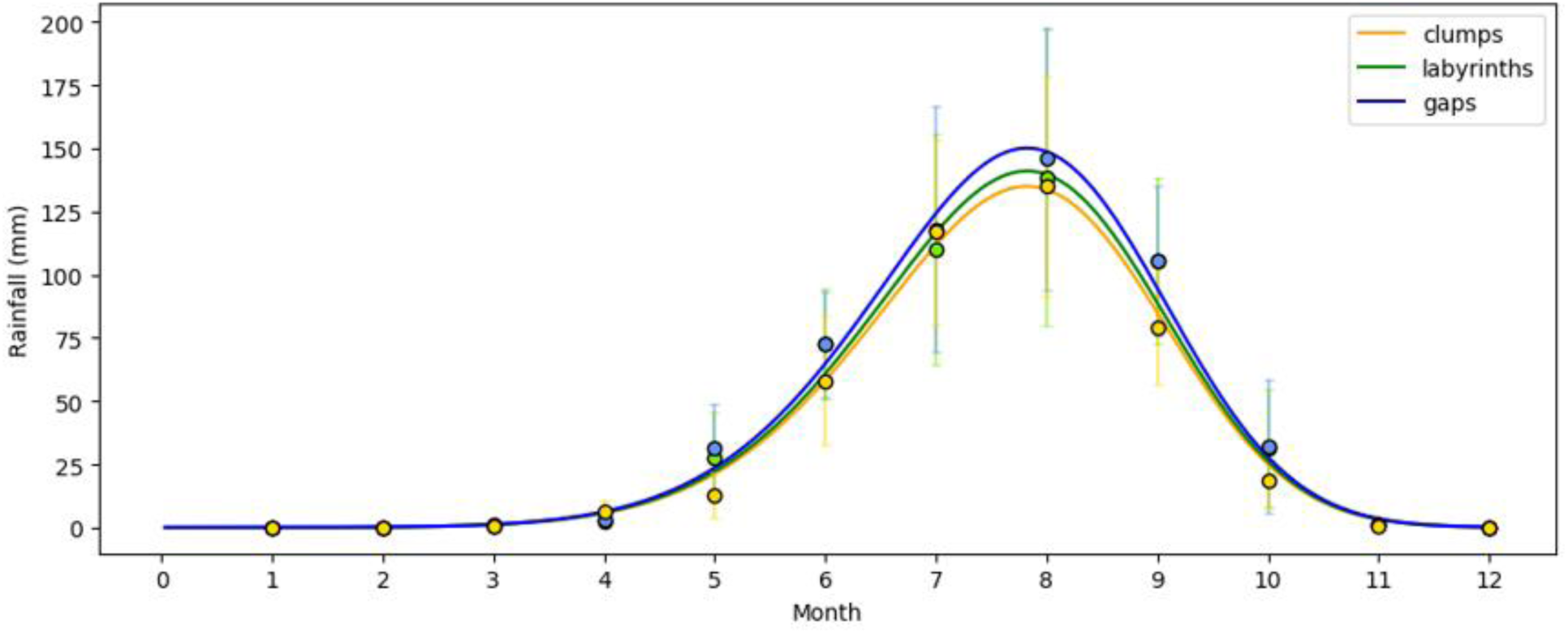
Rainfall function *R(M)* obtained with data collected at sites that show different types of patterns (clump, labyrinth, and gap). The dots represent medians across years, and error bars represent interquartile ranges. We fitted a skewed-gaussian function to represent seasonal precipitation. Fig. S2.2a-c shows the patterns observed in the regions where the rain data were collected; panels g-i show the simulated patterns obtained with these rainfall curves and added stochasticity.

Thus, we added a random number generated from a normal distribution with mean 0 and standard deviation 0.3*R(M),* the latter representing the observation that drier months show a lower rainfall variability (see Fig. S2.1). The resulting random rainfall values were distributed normally with mean *R(M)* and standard deviation 30% of the given rainfall.

We used these simulations with stochastic rainfall to check whether vegetation patterns changed seasonally or remained stable across seasons. As shown in Figure S2.2, both the stochastic rainfall simulations and fixed rainfall simulations exhibited similar patterns at similar annual rainfall levels, indicating that the vegetation patterns generated by this model are robust under stochastic and seasonal changes in rainfall. Therefore, for simplicity we used fixed rainfall values for our subsequent simulations and analyses.

**Figure S2.2:**
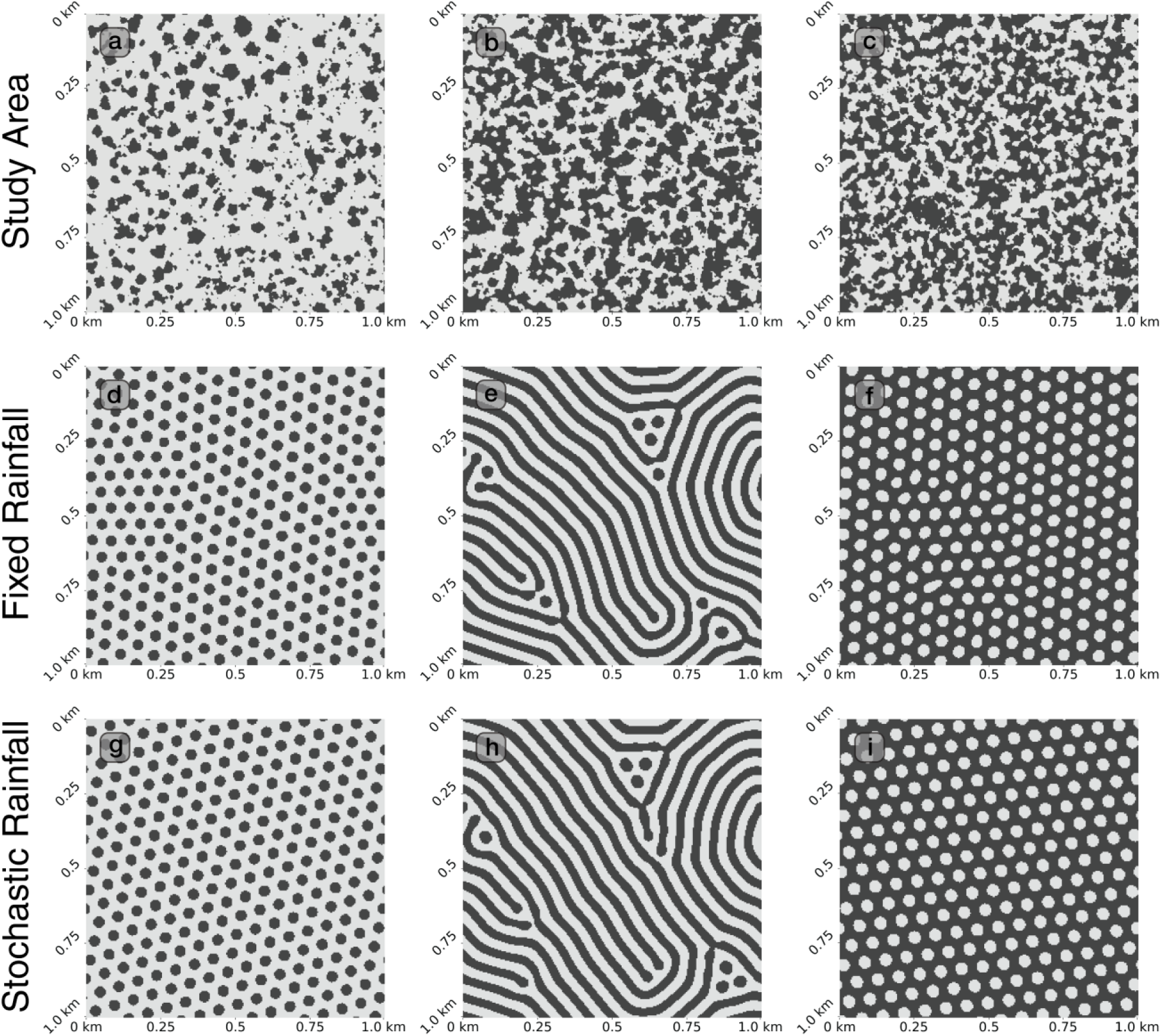
Simulated vegetation patterns and real patterns from our study area. Panels (a), (b) and (c) are satellite images derived from Google Earth pro, with annual precipitation 430.8, 478.0 and 508.9 mm/year, respectively. Panels (d), (e) and (f) correspond to simulations with fixed annual precipitation *r* = 430, 475 and 510 mm/year, respectively. Panels (g), (h) and (i) correspond to the stochastic rainfall simulations with annual precipitation *r* = 430, 475 and 510 mm/year, respectively. For all snapshots, black indicates the presence of vegetation and white means bare soil, with panels resulting from the binarization protocol explained in Section 4.

### Section 3. Model parametrization

For all model parameters, we used available information to constrain the range of possible values. Our main goal was to identify a plausible order of magnitude for each parameter. Whenever data were not available, we used theoretical calculations. In the absence of either option, which was the case for one parameter only, we resorted to sensitivity analyses to select a value that helped replicate the spatial patterns of our focal region.

#### 3a. Water-related parameters

##### Water diffusivity (Dw)

To calculate the diffusivity parameter, we first calculated the cross-sectional average velocity of water flow using the Manning equation (Manning, 1891):

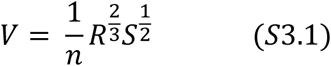

where *V* is the cross-sectional average velocity (m/s), *n* is Manning’s Roughness Coefficient for different surface material, *R* is the hydraulic radius (m, see Table S3.1), and *S* is the slope (elevation difference/section length)

**Table S3.1:**
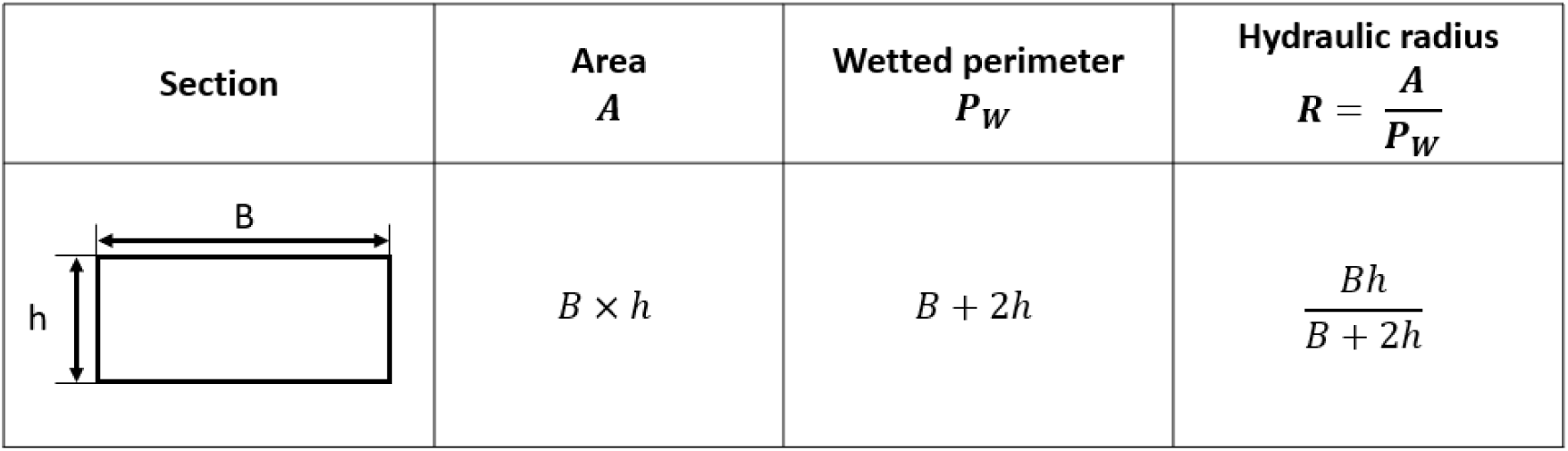
Channel section geometry and associated parameters.

| **Section** | **Area**<br>***A*** | **Wetted perimeter**<br>***P<sub>W</sub>*** | **Hydraulic radius**<br>$\boldsymbol{R = \frac{A}{P_W}}$ |
|---|---|---|---|
| B<br>h | *B* × *h* | *B* + 2*h* | $\frac{Bh}{B + 2h}$ |

To calculate the hydraulic radius, we assumed that the intense rainfall during the rainy season creates a very shallow sheet flow with width *B* = 1 m and depth *h* = 0.0001 m in a horizontal area with very small local slope (0-0.1%). These two values allowed us to calculate the hydraulic radius for a channel with a given rectangular cross-section (Table S3.1). For Manning’s roughness coefficient, we assumed that the surface material for the shallow sheet flow was similar to a clean and straight natural stream with no rifts or deep pools, which led to a value of 0.03 (Arcement and Schneider, 1989). With this information and a local slope of 0.05%, Eq. (S3.1) resulted in a flow velocity of 0.00161 m/s. Water diffusivity was determined by taking into account the flow velocity in all directions. Naively, because we assumed that *w* represents surface water (see main text), if water can travel 0.00161 m per second, then water moving in all directions will initially cover a surface π (0.00161)^2^ per second. The calculated value was thus 255 m^2^ per year (similar to the one used in Siteur et al. (2014)), which we considered as a maximum possible diffusivity since the area added per unit time is that of an annulus instead of a disc. With this in mind, we chose the more conservative *D_w_* = 170 m^2^ per year, a value within the same order of magnitude that also ensured numerical stability, is close to the one used in Klausmeier (1999) repeatedly adopted in theoretical work, and is in line with the value calculated in Ursino (2005).

##### Water loss rate (l)

To calculate the loss rate, we first calculated the potential evapotranspiration (PET) using the following equation (Thornthwaite, 1948):

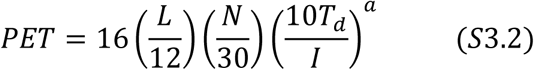

*PET* is the estimated potential evapotranspiration (mm/month), *T_d_* is the average daily temperature (degrees Celsius) of the month, *N* is the number of days in the month, *L* is the average day length (hours) of the month, *I* is a heat index which depends on the mean temperature, and the exponent *a* is given by the formula:

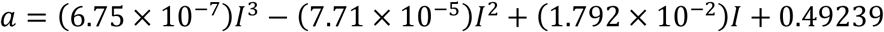

We set the averaged day length of the month (*L*) to 12 hours and the number of days in the month (*N*) to 30 days. For the rest of parameters, we used two cities within the study area as a reference, Al Nahud and Muglad, for which data were available. The average daily temperature (*T_d_*; Al Nahud: 27.8 ℃; Muglad: 28.5 ℃) and precipitation data (1960-1990) for these two cities (Al Nahud: 315 mm; Muglad: 471 mm) were derived from Alam et al. (2013). Based on the average daily temperature and the heat index table (Seed and Malik, 1968), the heat index *I* for Al Nahud was 13.49 and for Muglad was 13.94. Therefore, we could calculate the PET for each city and get the estimated water loss rate by dividing the annual PET to the mean annual precipitation. The estimated water loss rate range was from 3.7 to 5.4 *year*^−1^.

#### 3b. Plant-related parameters

##### Plant diffusivity (D_p_)

We interpreted the diffusivity parameter for plants (*D_p_*) as resulting from successful dispersal events for the focal species, considering germination rate and first-year seedling survival rate.

There is no literature indicating the specific seed dispersal strategy of *A. mellifera*, but it has been pointed out that the seeds of *A. mellifera* disperse over a relatively small distance near their parent plant (Donaldson, 1969). This means that the primary seed dispersal strategy can be barochory, and thus the seed dispersal of *A. mellifera* would primarily rely on gravity and be assisted by wind. The average monthly wind speed in the last two decades for our area of interest was under 5 m/s (FNMOC 10 m Surface Winds, erdlasFnWind10). Moreover, according to the Beaufort Wind Scale, the sample area typically experiences light to gentle breeze, which is unlikely to make a long-distance seed dispersal. Thus, we assumed that the seed dispersal distance outside the canopy equals the radius of the canopy (2.04 m, Figure S3.1) and calculated the circle of diffusion area based on the schematic diagram for the study of the effect of wind direction on seed dispersal (Hagos, 2001).

**Figure S3.1:**
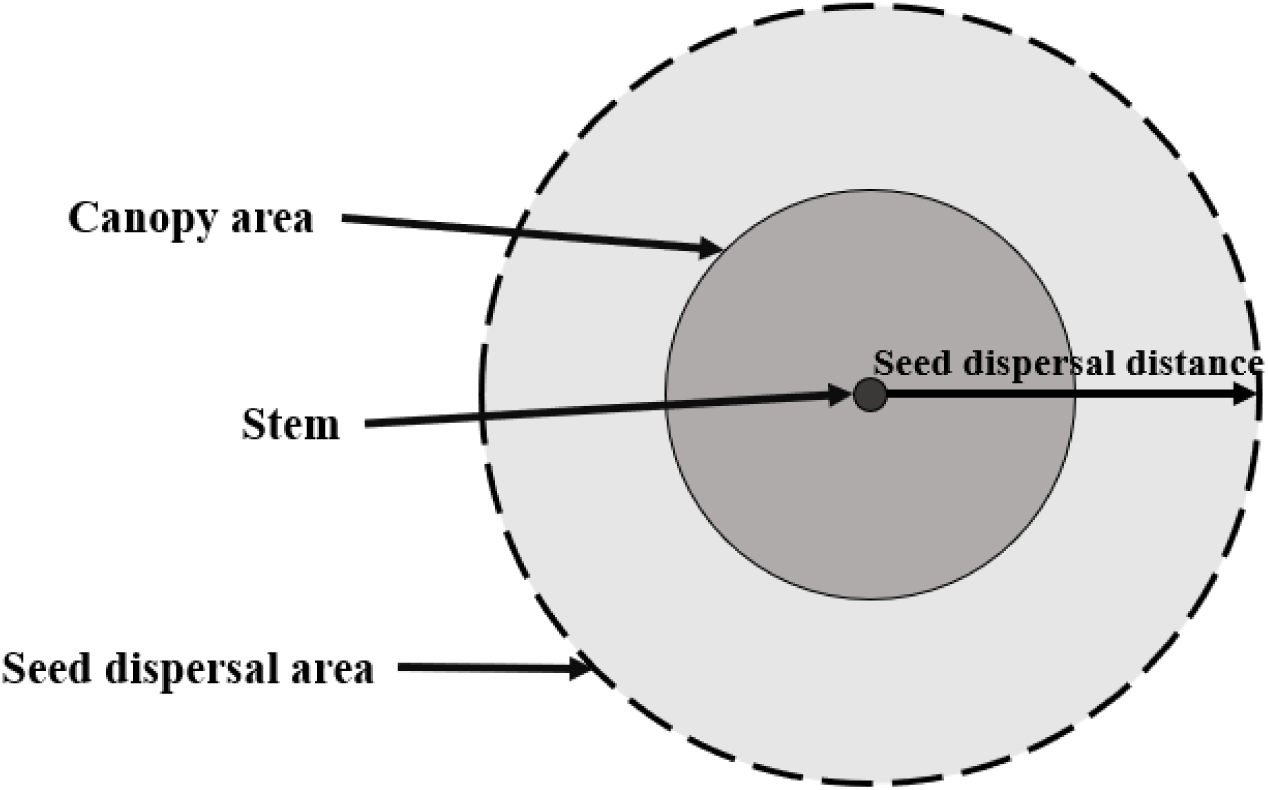
Schematic diagram showing *A. mellifera* seed dispersal. The annulus in light gray represents the area of successful seed dispersal.

Therefore, we considered the seeds that disperse outside the canopy as a potential successful dispersal event and assumed the dispersal radius to be twice that of the canopy, which leads to a dispersal area of 39.22 m^2^. We further weighted the success of dispersal by the survival probability after a year (and thus establishment probability). On one hand, 14% of *A. mellifera* seeds disperse outside the canopy (Hagos, 2001); on the other hand, the germination rate of *A. mellifera* seeds is around 50-80% (Bein et al., 1996), and the calculated first-year survival rate of seedlings outside the canopy is around 42% (Joubert et al., 2013). Altogether, the plant diffusivity calculated from the data above ranged 1.15-1.88 m^2^ per year, resulting from the following equation:

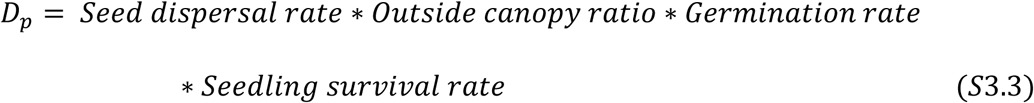

In addition, clonal reproduction of the shrub also contributes to its spatial spreading even in the absence of wind. To also account for this possibility, we used the more conservative value *D_p_* = 1 m^2^ per year, in line with the one used in Klausmeier (1999).

##### Other plant-related parameters

For the yield of plant biomass per unit water consumed (*y*) we used the assumptions made in the literature for encroacher species (*Terminalia sericea* and *A. mellifera*) of 1500 liters of water consumed to produce 1 kg dry mass (NAU, 2010), i.e. *y* = 0.00067 *kg_drymass_* per *kg_water_*.

In addition, two studies have mentioned the mortality rate of *A. mellifera*: Joubert et al. (2013) reported that around 1% of labeled seedlings survived after 6 years, which means the probability for *A. mellifera* seedlings to survive for 6 years is 0.01. Assuming that the survival probability does not change over the years and life stages, *A. mellifera* has a probability 0.46 to survive for one year, and therefore the annual mortality probability is 0.54. A later study showed that the mortality rate of shrubs over a 35-year period is around 61.3% (Joubert et al., 2017), which means that the probability to survive for one year is 0.97, and thus the annual mortality probability equals 0.03. These data provided a range of values for *A. mellifera* mortality probability (*m*) with 0.03 and 0.54 per year as extremes.

The parameter accompanying the water uptake term (*v*), on the other hand, has units that are not typical for an uptake rate ((*kg_drymass_m*^−2^)^−2^*year*^−1^), and therefore we could not find a value from available data or by following ecological arguments as above. A sensitivity analysis revealed that the main role of this parameter is to shift the rainfall values where patterns and desertification occur, but does not change the patterns. Due to the lack of information, we chose this parameter to ensure that (once the rest of parameters were fixed to realistic values, see above and Table S3.2), the range of rainfall values for which patterns emerged was comparable to those observed in the focal system. Thus, we used a value of *v = 57*, but other values of *v* that we tried (*v = 25, 50, 60, 75, 100*) did not alter our conclusions qualitatively.

**Table S3.2:** List of model parameters, values and units. For parameter values or ranges directly obtained from the literature, we included the corresponding reference; for values that we calculated based on ecological and/or physical arguments, we included the section with the corresponding explanation of the steps.

| Parameters | Description | Units | Calculated range | Simulation value | Reference |
|---|---|---|---|---|---|
| *w* | Water density | *kg*<sub>*H2O*</sub>*m*<sup>−2</sup> | variable | variable | N/A |
| *p* | Plant density | *kgm*<sup>−2</sup> | variable | variable | N/A |
| *r* | Precipitation rate | *kg*<sub>*H2O*</sub>*m*<sup>−2</sup>*year*<sup>−1</sup> | 430-509 | 380-540 | CHIRP precipitation database |
| *l* | Water loss rate | *year*<sup>−1</sup> | 3.7-5.4 | 5 | Section 3a; Eq. S3.2 |
| *v* | Water uptake rate | (*kg*<sub>*drymass*</sub>*m*<sup>−2</sup>)<sup>−2</sup>*year*<sup>−1</sup> | - | 57 | - |
| *y* | Yield of plant biomass per unit water consumed | *kg*<sub>*drymass*</sub>(*kg*<sub>*H2O*</sub>)<sup>−1</sup> | 0.00067 | 0.00067 | NAU, 2010 |
| *m* | Plant mortality rate | *year*<sup>−1</sup> | 0.03-0.54 | 0.5 | Joubert et al., 2013; Joubert et al., 2017 |
| *D*<sub>*w*</sub> | Diffusion coefficient for water | *m*<sup>2</sup>*year*<sup>−1</sup> | 255 (max) | 170 | Section 3a; Eq. S3.1 |
| *D*<sub>*p*</sub> | Diffusion coefficient for plants | *m*<sup>2</sup>*year*<sup>−1</sup> | 1.15-1.88 | 1 | Section 3b; Eq. S3.3 |
| *t* | Time | *year* | variable | variable | N/A |
| *dt* | Time resolution | *year* | - | 0.001 | N/A |
| *dx* | Spatial resolution | *m*eter | - | 4 | N/A |
| *dN* | Perturbation | *kgm*<sup>−2</sup> | - | 1.0/0.1 | N/A |

### Section 4. Processing of vegetation images through unsupervised classification and clump detection

We analyzed snapshots from our simulations as well as 1km x 1km optical satellite images with 1.2 m spatial resolution for the focal study area extracted from Google Earth Pro. Note that Google Earth Pro provides only visible-band composites, so no near-infrared channel was available and vegetation indices such as NDVI or EVI could not be computed for these scenes. Thus, given the lack of information to map image colors into vegetation density and following standard practice, we binarized images before analyzing them. Here, in order to detect and discriminate vegetation from the surrounding bare soil, we used an unsupervised classification algorithm known as KMeans (Arthur and Vassilvitskii, 2007) that is widely used in image analysis, particularly in remote sensing applications for land cover classification, change detection, and vegetation analysis (Celik, 2009; Olaode et al., 2014; Ahmed and Akter, 2017; Wan et al., 2018).

KMeans clustering involves the partition of a dataset into *k* optimal distinct clusters, through the iterative minimization of a function *ψ*. More specifically, when provided an integral input *k* on a dataset *χ* ∈ ℝ*^d^*, the KMeans algorithm calculates *k* cluster centers *C* to minimize the sum of Euclidean distances from each point in the dataset to its nearest cluster center. Mathematically, this is achieved by minimizing the following function (Arthur and Vassilvitskii, 2007):

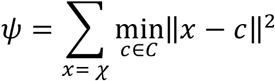

Because our aim is to binarize images, in our case we applied a binary KMeans classification (*k* = 2) to our satellite and simulated images, which resulted in a classification of image pixels into two clusters, as either “low” intensity pixels (representing bare soil) or “high” intensity pixels (representing vegetation) (see example in Fig S4.1). For our analyses, we used the KMeans implementation from the scikit-learn Python library (sklearn.cluster.KMeans), which features several enhancements over naïve KMeans. Specifically, we used the KMeans++ initialization strategy, which improves clustering performance and convergence stability by selecting initial cluster centers that are probabilistically distant from one another (Arthur and Vassilvitskii, 2007). Because simulation snapshots were essentially single-channel plant-density data, they were classified directly on that intensity, which is a standard step in digital image processing (Zhao et al., 2019); on the other hand, we classified satellite images on the full three-channel RGB values, therefore retaining the colour contrast between canopy and bare soil rather than discarding it by converting to greyscale.

Once images were binarized, we proceeded to identify and quantify the motifs, i.e. vegetation clumps. Thus, we used the two-pass Hoshen–Kopelman algorithm from the SciPy Python library (scipy.ndimage.label) to label individual classified vegetation clumps as connected components (Hoshen and Kopelman, 1976). While this labelling was sufficient to estimate the number of clumps, estimating clump sizes required additional care due to the potential presence of transient adjoining clumps in analyzed snapshots. These transient motifs occur when the pattern is close to but not yet at its full stationary state, and results from clumps still being in the process of forming/separating. Such artifacts affect to some extent the average clump size, and most certainly the maximum clump size measured with the snapshot. To eliminate these artifacts, we used an unsupervised Isolation Forest (Liu et al., 2008) applied to eight morphological discriminants/properties per clump (see table S4), the most diagnostic being the depth of constrictions in the width profile along the clump’s medial axis (Zhang and Suen, 1984), followed by distance-transform peak topology (Maurer et al., 2003) and a composite elongation indicator (Walton, 1948). Flagged clumps were isolated one to three levels earlier in the isolation trees than retained clumps (against a maximum depth of eight) and carried markedly lower anomaly scores, confirming that the flag was valid. This additional screening allowed us to reliably calculate average and maximum clump size.

As a side note, KMeans (similarly to other clustering algorithms) is sensitive to the choice of *k*: higher choices often lead to the detection of more features, but at the risk of overfitting and loss of statistical power. To ensure that our choice of *k = 2* was appropriate, we performed a silhouette analysis (Rousseeuw, 1987) and calculated the percentage of unexplained variance for analyses using k ranging from 2 to 10 (not shown). For both the satellite data as well as the simulated data, we found that *k* = 2 was the most parsimonious choice.

**Fig S.4.1:**
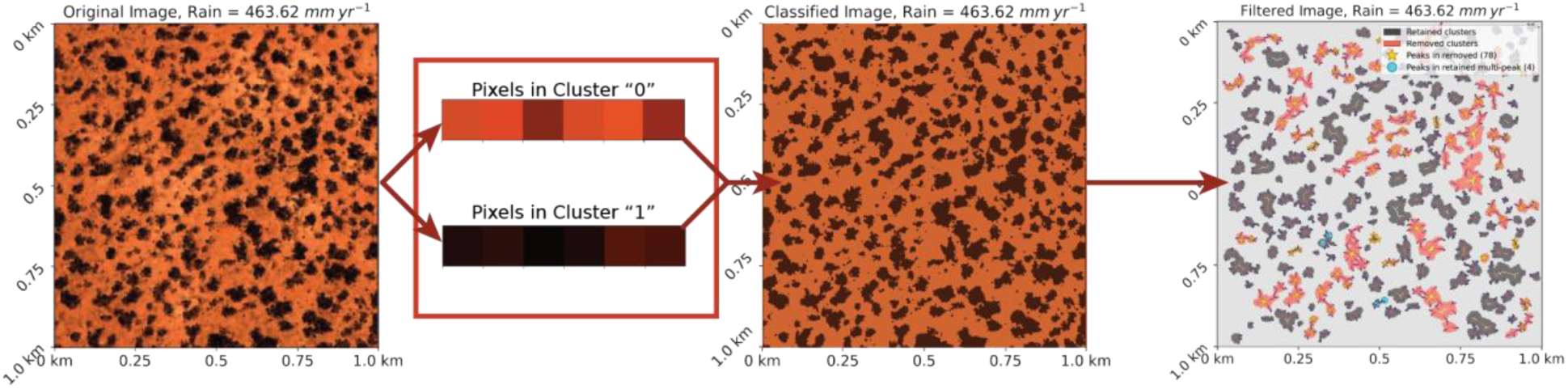
Example of workflow steps for the processing of a snapshot from our study area (left panel), which we first binarized version using the KMeans++ algorithm (center) and then screened to remove motifs that result from merged clumps (right).

**Table S4:**
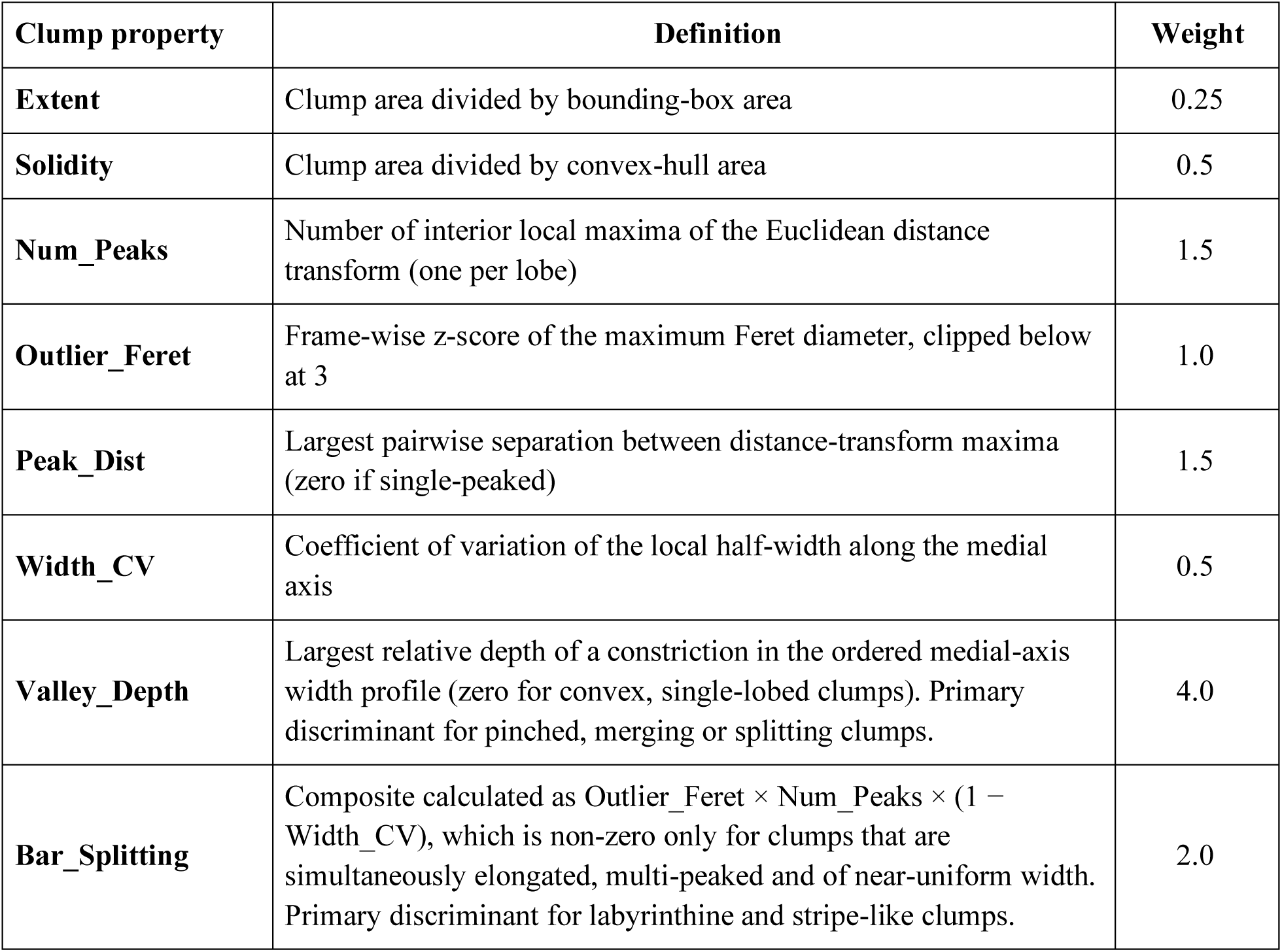
List of the eight morphological discriminants supplied to the Isolation Forest, with their respective weights.

| **Clump property** | **Definition** | **Weight** |
|---|---|---|
| **Extent** | Clump area divided by bounding-box area | 0.25 |
| **Solidity** | Clump area divided by convex-hull area | 0.5 |
| **Num_Peaks** | Number of interior local maxima of the Euclidean distance transform (one per lobe) | 1.5 |
| **Outlier_Feret** | Frame-wise z-score of the maximum Feret diameter, clipped below at 3 | 1.0 |
| **Peak_Dist** | Largest pairwise separation between distance-transform maxima (zero if single-peaked) | 1.5 |
| **Width_CV** | Coefficient of variation of the local half-width along the medial axis | 0.5 |
| **Valley_Depth** | Largest relative depth of a constriction in the ordered medial-axis width profile (zero for convex, single-lobed clumps). Primary discriminant for pinched, merging or splitting clumps. | 4.0 |
| **Bar_Splitting** | Composite calculated as Outlier_Feret × Num_Peaks × (1 − Width_CV), which is non-zero only for clumps that are simultaneously elongated, multi-peaked and of near-uniform width. Primary discriminant for labyrinthine and stripe-like clumps. | 2.0 |

### Section 5. Fourier transform and Power Spectrum

A Fourier transform shows one peak for every direction where there is periodicity in the corresponding image. The location of the peak represents how many times the motif is repeated within the picture (spatial frequency). For example, for the clump pattern (Fig. S5.1a), the Fourier transform (Fig. S5.1b) shows a peak at a distance from the origin that reflects the number of times that a clump is found in a given direction. Thus, tracing parallel lines that cross clumps in a given direction, the distance to the origin of each Fourier transform peak corresponds with the number of lines with such orientation within the picture; the orientation determines the angle at which the peak is located in the Fourier transform. The radial power spectrum integrates the Fourier transform across all possible angles and, in consequence, only depends on the distance to the origin (i.e. spatial frequency). Putting all together, any links connecting first/nearest neighbors in the original pattern (e.g. pink and brown lines in Fig. S5.1a) lead to peaks of the Fourier transform at the shortest possible distance to the origin (pink and brown circles in Fig. S5.1b) and contribute to the same peak in the radial power spectrum (brown solid box in Fig. S5.1c). This peak is the so-called fundamental peak, which determines the main (spatial) frequency within the picture. Because the fundamental peak provides information about the typical distance to nearest neighbors, its presence and location are used to characterize regularity within images/patterns (Deblauwe et al., 2011). Theoretically, this peak (and its harmonic peaks, located at twice, thrice, etc. the distance of the first peak) are typically the focus of research.

The clump and gap patterns, however, also show an additional and typically ignored feature. When considering lines that connect secondary neighboring clumps (a.k.a. next-nearest neighbors, green and red lines in Fig. S5.1a), because of the overlap of one row and the following one, the distance between lines is smaller, i.e. there are more lines than for first neighbors. These lines lead to farther peaks in the Fourier transform (red and green circles in Fig. S5.1b), and thus produce a secondary peak in the power spectrum (dark red box in Figure S5.1c) that is close to, but different from, the second harmonic of the fundamental peak (dashed-line box). Technically, this peak is a linear combination of modes (e.g. in Fig. S5.1b, the coordinates of the red circle can be obtained by adding the vectors of peaks from the first neighbors, specifically twice the vector of the purple circle plus the vector symmetric to the brown circle). Moreover, symmetry arguments place this peak at a location in the Fourier space (i.e. spatial frequency) that is exactly 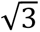 times the location of the primary fundamental peak in an ideal hexagonal pattern: since next-nearest neighbors are separated by the same distance as the flat-to-flat distance in a hexagon, calling *a* the distance between nearest neighbors (identical to the side length of the hexagon), then the distance between next-nearest neighbors is 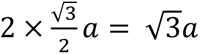, and thus so are their spatial frequencies.

**Figure S5.1:**
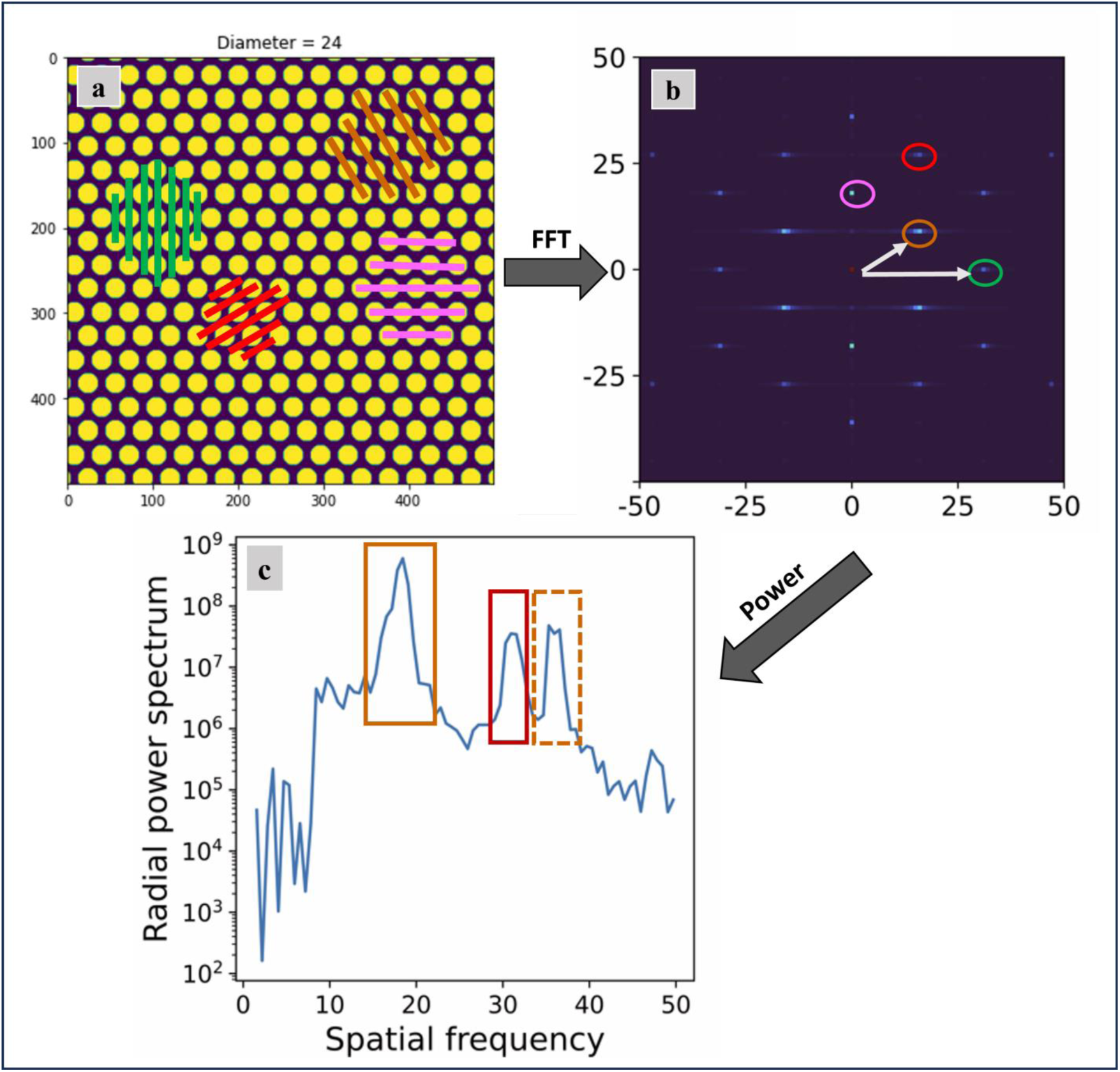
Link between pattern and power spectrum for an idealized (i.e., perfectly hexagonal) clump pattern. Panel (a) is the original image with red, pink, brown, and green lines indicating directions/lines crossing the center of the clump, the number of lines corresponds to spatial frequency in panels (b) and (c). Panel (b) shows the corresponding 2D Fast Fourier Transform (FFT), with dark blue indicating lower values and light blue indicating higher magnitude values. The horizontal and vertical axes correspond to the spatial frequencies in the horizontal and vertical directions respectively, and the color of circles corresponds to those of the lines in panel (a). Panel (c) shows the radially averaged power spectrum calculated from the 2D FFT. The primary fundamental peak, denoted by a brown solid box, represents the spatial frequency of the brown and pink lines in panel (a) (and circles in panel (b)), while its harmonic peak is identified by the dashed brown box. The secondary fundamental peak, indicated by the red box, corresponds to the spatial frequency of the red and green lines in panel (a) (circles in panel (b)).

#### 5.1 Relationship between power spectrum and pattern morphology

To investigate the behavior of this secondary fundamental peak, we constructed perfectly hexagonal clump patterns and first varied the size of the clumps only (i.e., number and position of clumps remained constant, see Fig. S5.2). As expected, because the location of the clumps remains fixed, the location and height of the primary fundamental peak in the power spectrum did not change, and neither did the location of the secondary fundamental peak (see above); however, the height of the secondary fundamental peak changed non-monotonically: it decreased with clump size until clump size was ∼20 pixels, then started increasing again. At that size threshold, the harmonic of the primary fundamental peak started decreasing. These changes corresponded with changes in the intensity of the second line of peaks in the Fourier transform (see red boxes for the secondary fundamental peak and green boxes for the harmonic of the primary fundamental peak).

**Figure S5.2:**
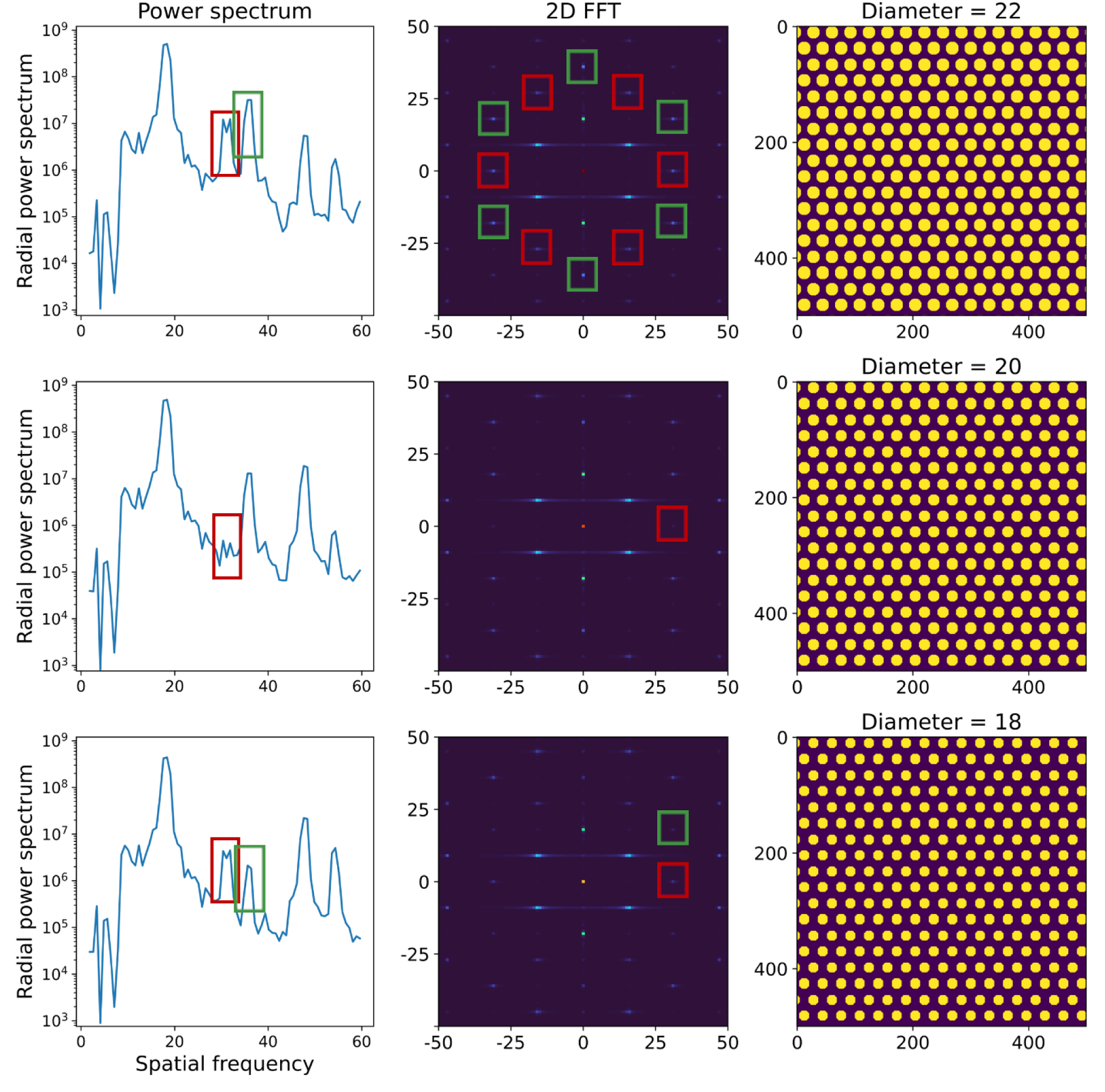
Power spectra (left) and Fourier transform (center) for idealized clump patterns (right) with identical number and location, but different sizes (clump diameter changes from 22 to 18 pixels from top to bottom). The peaks in the power spectrum and associated peaks in the Fourier transform associated with the secondary fundamental peak are denoted by red boxes, and those associate with the harmonic of the primary fundamental peak by green boxes.

To better understand these changes in the height of the secondary fundamental peak, we then reduced the number of clumps and monitored the power spectrum as a function of both clump size and number (Figure S5.3-S5.5). The power spectrum obtained with just five clumps showed now a well instead of a peak, with the location of the well increasing as clump size decreased and the depth of the well being slightly affected by size changes (Fig. S5.3). We then removed four of the clumps to leave the central one, which produced a well at the same spatial frequency as that for the five clumps (Figure S5.4), with a location that also increased as clump size decreased but a depth that barely changed. Finally, when we instead multiplied by nine the number of clumps (i.e. 45 clumps in total, Figure S5.5), the depth and position of the well was much more sensitive to clump size: as size decreased, the well moved toward higher spatial frequencies, leading to the decrease and eventual disappearance of the secondary fundamental peak for intermediate clump sizes, then its reappearance and decrease of the harmonic of the primary fundamental peak as clump size is decreased further.

Taking all the above into consideration, we conclude that changing size and number in such patterns leads to changes in the location of the well and secondary fundamental peak and how much they overlap (and therefore how much they offset each other); the conclusion is also applicable (albeit for smaller clump sizes) to the harmonic of the primary fundamental peak. This explains why the heights of the secondary fundamental peak and the harmonic change, and point out that motif size and number are important features to monitor when analyzing vegetation patterns such as clumps and gaps. The effective behavior of these two peaks can also be approached from a more qualitative perspective: as clumps (or gaps) converge to labyrinths, distances between neighbors become more permanent while next-nearest neighbors cannot be defined anymore, that is, the primary fundamental peak remains but the secondary fundamental peak ultimately disappears.

**Figure S5.3:**
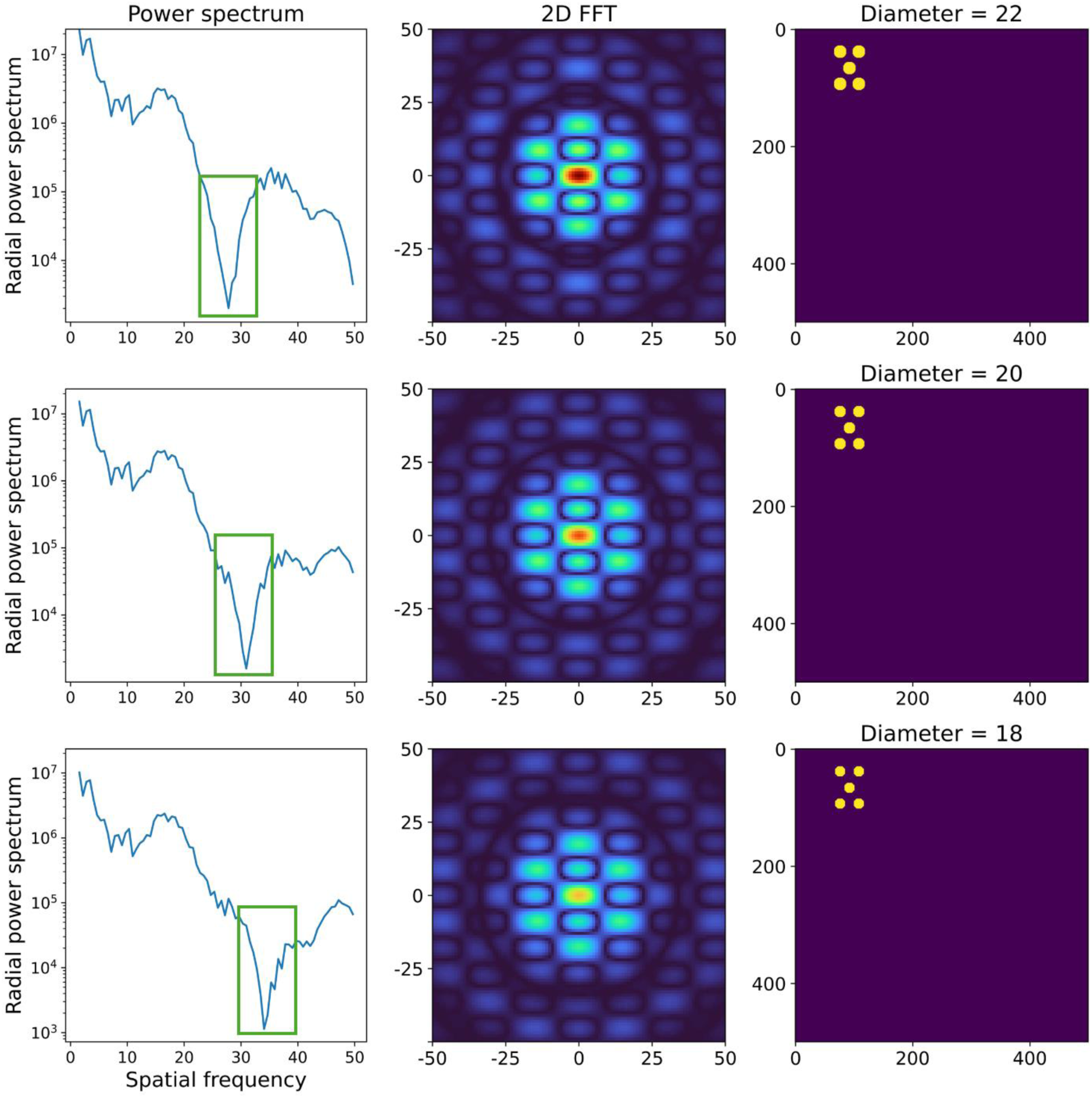
Power spectra (left) and Fourier transform (center) for idealized clump patterns (right) with identical number and location, using only five clumps. Note that the secondary fundamental peak is not present, but instead a well between the first peak and its harmonic emerges (see green box).

**Figure S5.4:**
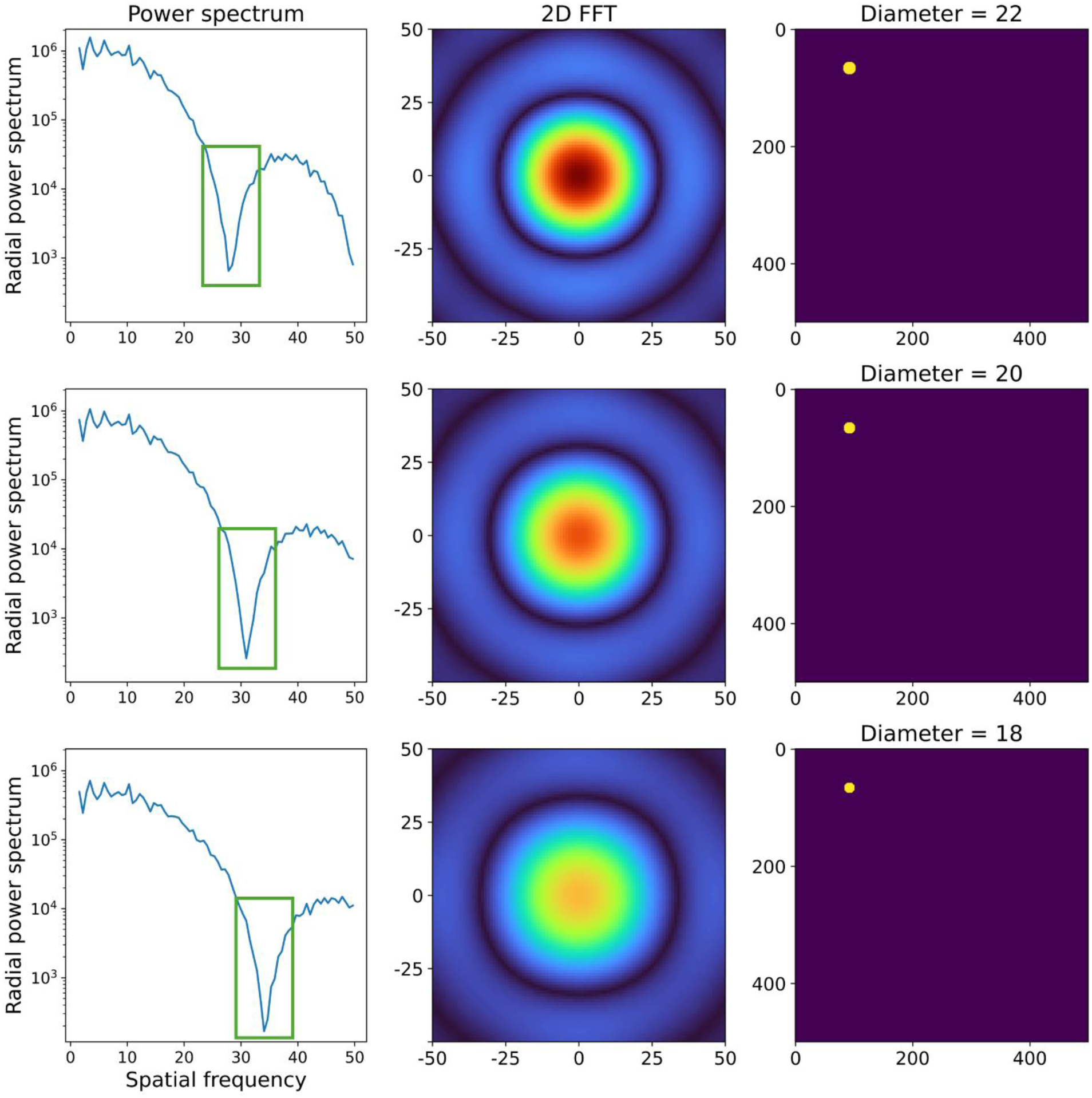
Same as in Fig. S5.3 but using a single clump in an otherwise empty system.

**Figure S5.5:**
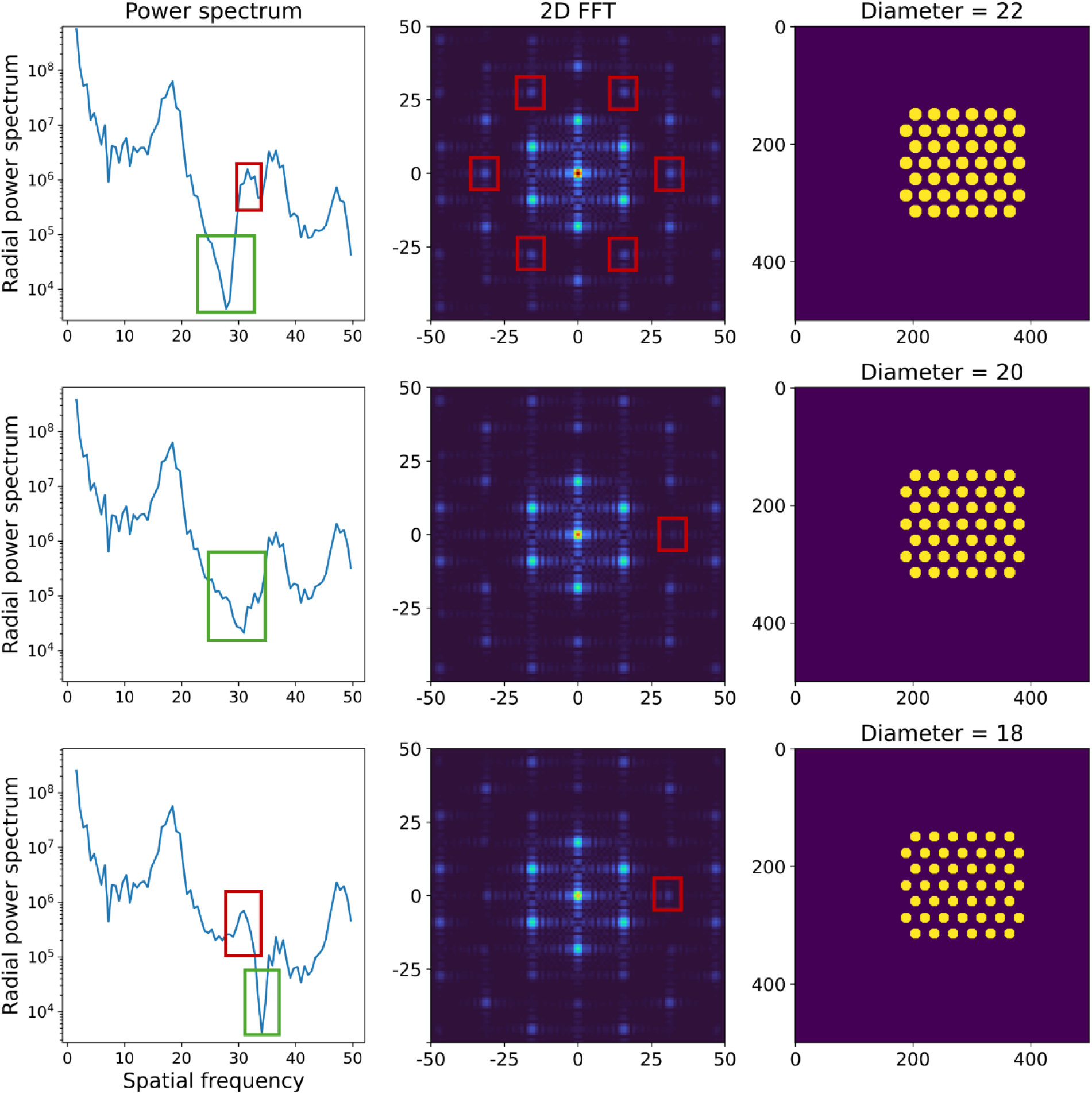
Power spectra (left) and Fourier transform (center) for idealized clump patterns (right) with identical number and location, with 45 clumps. The secondary fundamental peak is present in the power spectrum and marked with a red box, as well as the associated peaks in the Fourier transform. Note that, for the case with 20-pixel-diameter clumps (middle row), both the secondary fundamental peak and the well overlap, resulting in a shallower well in the power spectrum and lower intensity in the peaks of the Fourier transform (see red boxes).

#### 5.2 Busse balloon and Eckhaus instabilities for the Klausmeier model

To understand better the behavior of the primary and secondary fundamental peaks as rainfall changes, we extended the typical study of stability of regular patterns. Classically, the study of these patterns entails the identification of the regions of the phase diagram (here, the values of rainfall) for which the stationary state of the non-spatial system (i.e. the mean-field case) is stable, but the spatial system is unstable to perturbations (Klausmeier, 1999). These are the so-called Turing-unstable regions (Turing, 1952). The calculation was later extended for a one-dimensional version of the modified Klausmeier model to consider additional regions where the spatial system may form stable regular patterns after being perturbed: the so-called Busse balloon (e.g. Siteur et al., 2014). Delimited by lines called Eckhaus instabilities where the pattern is marginally stable, the Busse balloon is a region of the phase space within what would be the desertic phase in the mean-field case; these are regions that, despite not being Turing-unstable, regular pattern can emerge. The Busse balloon thus defines the maximum theoretical range for spatial bistability, although whether patterns actually emerge (i.e. the realized spatial bistability range) depends on how the system was initially perturbed. It has been argued that this pattern multistability offers the system various ways to respond to aridity, potentially leading to a smooth instead of an abrupt transition (Bastiaansen et al., 2018), although the generality of this hypothesized mechanism has been recently contested (Pinto-Ramos & Martínez-García, 2026).

Aiming to understand our observed phenomenology, i.e. an abrupt transition preceded by constant clump size and decreasing number of clumps, we used our own calculation of the Eckhaus instabilities for our specific model and parametrization (see technical details in Halder and Bonachela (2026)). In a nutshell, we aimed to determine the range of pattern wavelengths that could emerge for a given rainfall, which required assessing the stability of each periodic solution to disturbances that slowly modulate the spacing of the pattern itself (i.e. the Eckhaus mechanism (Eckhaus, 1965)). To this end, we computed families of stationary periodic wavetrain solutions for our focal model (Eqs.(1)-(2)) in one spatial dimension across the rainfall gradient, and evaluated the stability of each solution by formulating and solving the associated discrete Bloch operator problem (Kittel and McEuen 2015; Sherratt 2013).

As Fig. S5.6 shows, the wavenumber of the patterns generated by our two-dimensional simulations fall within the regions of pattern stability of the one-dimensional Eckhaus instability calculations (dashed grey lines). A simple symmetry argument can explain why: the emergent patterns within this region are all very close to a hexagonal pattern, which is the result of alternating rows of clumps shifted a certain distance from each other (specifically, if the distance between clumps within a row is *a*, rows are shifted a distance *a/2* from each other); in other words, the two-dimensional pattern is composed by a collection of one-dimensional ones that set the main distance to nearest neighbors and, therefore, the wavenumber of the pattern (see pink and brown lines in Fig. S5.1).

Moreover, similar symmetry arguments can help relate the number of clumps with this main wavenumber of the pattern (see Fig. S5.6). Assuming again the pattern to be hexagonal (which is the idealization of the clump patterns in our system), one can calculate the total area of the system as the area of the triangles that form each hexagon multiplied by the number of triangles in the system. Given that each triangle is formed by one clump in a row and two in the next row, the total number of triangles is approximately 2×*n_clumps_*. Calling the distance between nearest neighbors *a* (inversely proportional to the wavenumber), the area of each triangle is 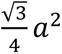*a*^2^. Finally, in a two-dimensional system of linear size *L*, the total area of the system is *L^2^*. All together, leads to the following relationship between the wavenumber of the pattern K and the number of clumps:

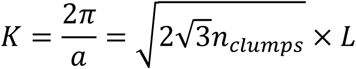

This approximation becomes less exact as clump patterns depart from ideal hexagonal patterns. Nonetheless, with this relationship in mind, Fig. S5.6 provides an explanation as to why the number of clumps decreases with aridity: for low rainfall levels, the number of clumps is the largest that it can be in a stable regular pattern, and thus the wavenumber follows closely the upper Eckhaus instability line, which shows a negative slope as rainfall decreases.

**Figure S5.6:**
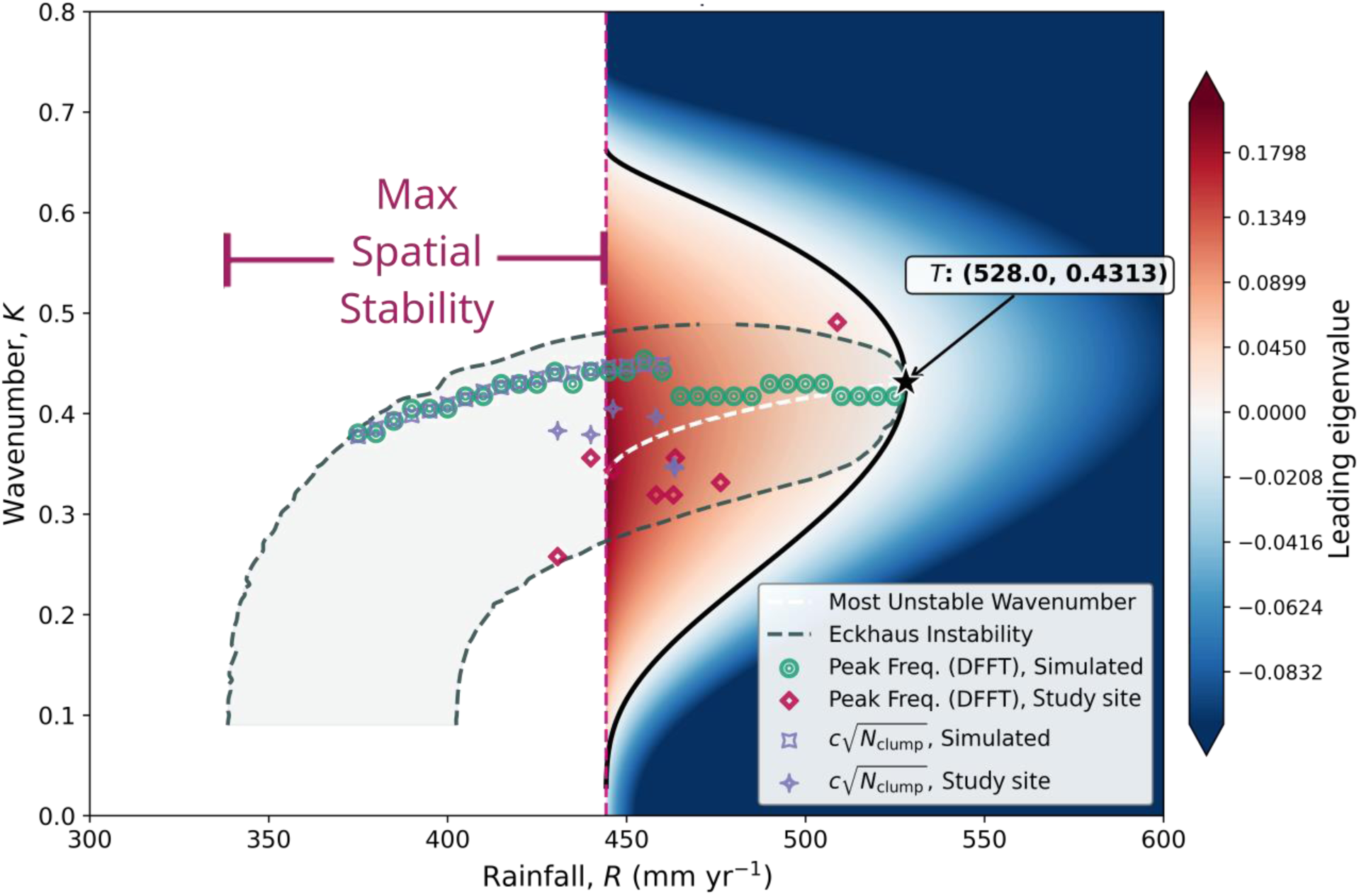
Theoretical calculation of where patterns emerge for the one-dimensional generalized Klausmeier model, overlapped with results from our two-dimensional simulations and the analyzed satellite pictures. The region delimited by the vertical dashed line and the solid black line represents the region where the mean-field system is stable but the complete spatial model is unstable and thus patterns emerge after using perturbations of wavenumber *K* (colors represent sign of the leading eigenvalue, and thus stability to these perturbations, with red indicating that patterns emerge and blue that they do not). This is the classic “Turing-pattern” zone, for which the lower R limit coincides with that of mean-field bistability. Dashed grey lines indicate the Eckhaus instabilities (marginally stable periodic solutions), with shaded area representing where stable patterned solutions can emerge (i.e. Busse balloon). As justified above, the region between the vertical line (beginning of mean-field bistability) and the smallest rainfall for which the Eckhaus instability is defined represents the maximum range of R values for which spatial bistability emerges. Green dots and red diamonds represent the wavenumber of the primary fundamental peak for simulations and real system, respectively. Grey symbols represent the approximated wavenumber of the primary fundamental peak, calculated as the square root of the number of clumps in the system, scaled by a given factor (see text above)

#### 5.3 Power spectrum for images from the focal area

We calculated the power spectrum associated with images from our study area, to explore whether they would allow us to identify a changing secondary peak and therefore the bistability region. Unfortunately, the secondary fundamental peak was not discernible due to the heterogeneity and stochasticity associated with real patterns and environmental conditions (and/or, as explained in main text, because the rainfall values explored correspond to values for which the tailored Klausmeier predicts the secondary peak to be negligible, see Fig. S5.7, which shows the radial-averaged power spectrum of clump patterns with rainfall levels ranging from 430.8 to 463.6 mm/year). Nonetheless, placing the wavenumber of the patterns in the Busse balloon of our Klausmeier model with data-informed parameterization (Fig. S5.6) shows that the remote-sensing patterns can be explained by this region. Moreover, it allows us to confirm the decreasing trend in the number of clumps as aridity increases (not fully clear in Fig. 6), since the shape of the Busse balloon forces the eventual decline as rainfall decreases.

**Figure S5.7:**
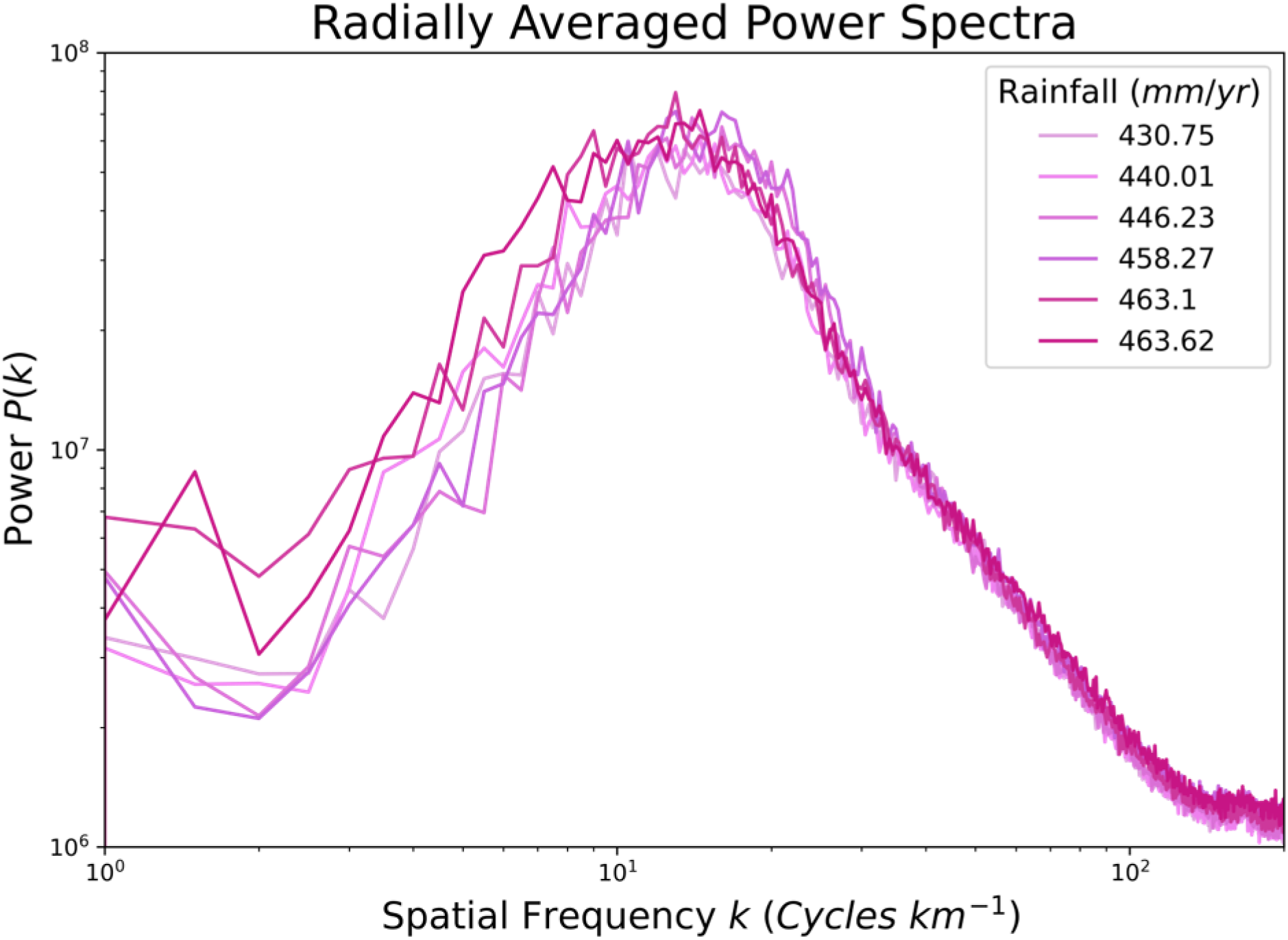
Radially averaged power spectrum of the focal area’s clump patterns observed for rainfall ranging from 430.8 to 463.6 mm/year.

### Section 6. Results with other model for vegetation patterns in semi-arid ecosystems

To investigate which features of our results are specifically linked to bistability, we repeated the analysis with other well-accepted models for vegetation in semi-arid ecosystems. In particular, we sought to explore all possible combinations of bistability. As explained in the main text, here we differentiate between stability for both vegetated and desertic states when space is not accounted for (mean-field bistability), and when space and patterns are present (spatial bistability), and the distinction is not only technical but also of ecological importance. The existence of non-spatial bistability indicates that the ecological interactions present in the system (in this case, local competition and facilitation) allow for the coexistence of both vegetated and desertic states, with the realized state depending on past history of the system. Spatial bistability (in this case, accompanied of spatial patterns) indicates that the fact that these interactions also occur at different scales is important not only for the generation of the patterns but also to maintain that coexistence.

Thus, we simulated i) a model that shows patterns but no bistability (i.e. shows a smooth transition in both non-spatial and spatial versions, (Martinez-Garcia et al. 2013)); ii) a model with only spatial bistability (that is, the non-spatial version of the model shows a smooth transition, but the spatial version shows bistability, (Rietkerk et al. 2002)); and iii) a model that, like the Klausmeier model, shows both non-spatial and spatial bistability (Siero, 2020). We used generic parametrizations for these models, as we lacked information to tailor their parameter values to our study area.

#### i) Model without spatial nor non-spatial bistability

The model in Martinez-Garcia et al. (2013) replicates all three vegetation patterns observed in semi-arid ecosystems. Differently from the Klausmeier model, however, patterns emerge purely from vegetation growth and competitive interactions (i.e. it does not implement vegetation biomass, but the equation encodes interaction between plants through a non-local competition kernel that affects vegetation growth rate within a constant interaction radius. Rainfall or aridity are not directly included in the model, but rather indirectly represented through the maximum growth rate, which is the parameter whose change triggers desertification or pattern changes. There is only one equation, which represents a unique value of the average vegetation density expected for each maximum growth rate (see Fig.S6.1). In other words, the desertification transition is smooth and there is no spatial bistability when the system approaches the transition.

Using the parametrization proposed in (Martinez-Garcia et al. 2013), which was inspired by vegetation in semi-arid ecosystems, we obtained the power spectra depicted in Fig. S6.2. Similarly to our tailored Klausmeier model, there is a secondary peak next to the harmonic, and both change heights as growth conditions change. Differently from the Klausmeier model, however, the location and height of the primary fundamental peak did not change. All together indicates that clumps changed in size but not in location (and therefore not in number, see Section 5 above). Indeed, Figs. S6.3 shows that both average and maximum clump size decreased with worsening growth conditions (lower maximum growth rate parameter, *β*) whereas the number of clumps remained approximately constant. This behavior contrasts with that observed in the spatial bistability region of the Klausmeier model, but agrees with the behavior observed outside of that spatial bistability region. Therefore, these results confirm that spatial bistability of the emergent patterns drives the decrease in the number of clumps, approximately constant in size, observed in the Klausmeier model and in the study area as aridity increases.

**Figure S6.1:**
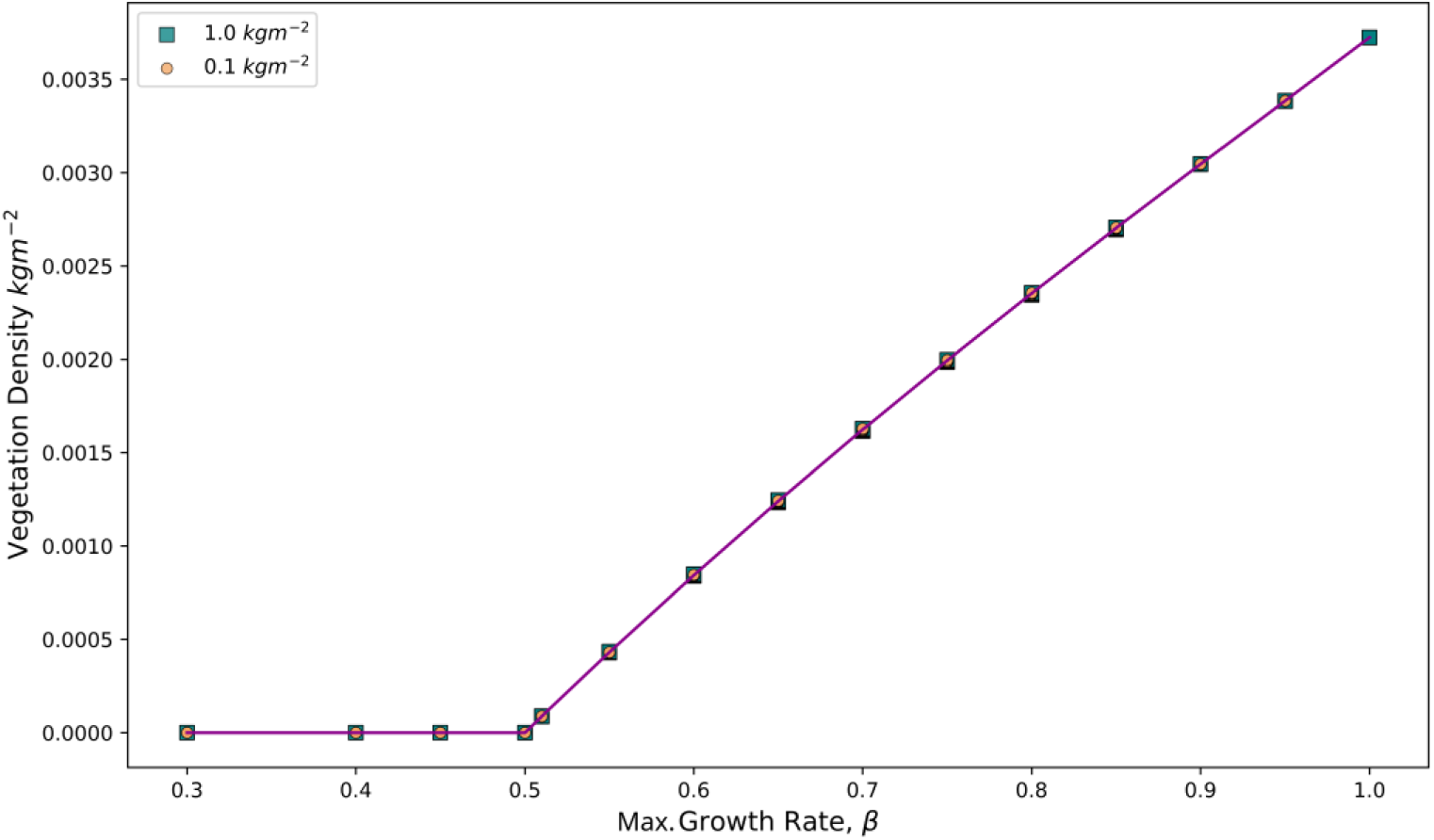
Phase diagram for the model in (Martinez-Garcia et al. 2013). The desertification transition, which occurs at a maximum growth rate 0.5 is smooth and therefore there is no bistability; only one average vegetation density is possible for every value of the maximum growth rate, which makes this transition much more predictable than an abrupt one.

**Figure S6.2:**
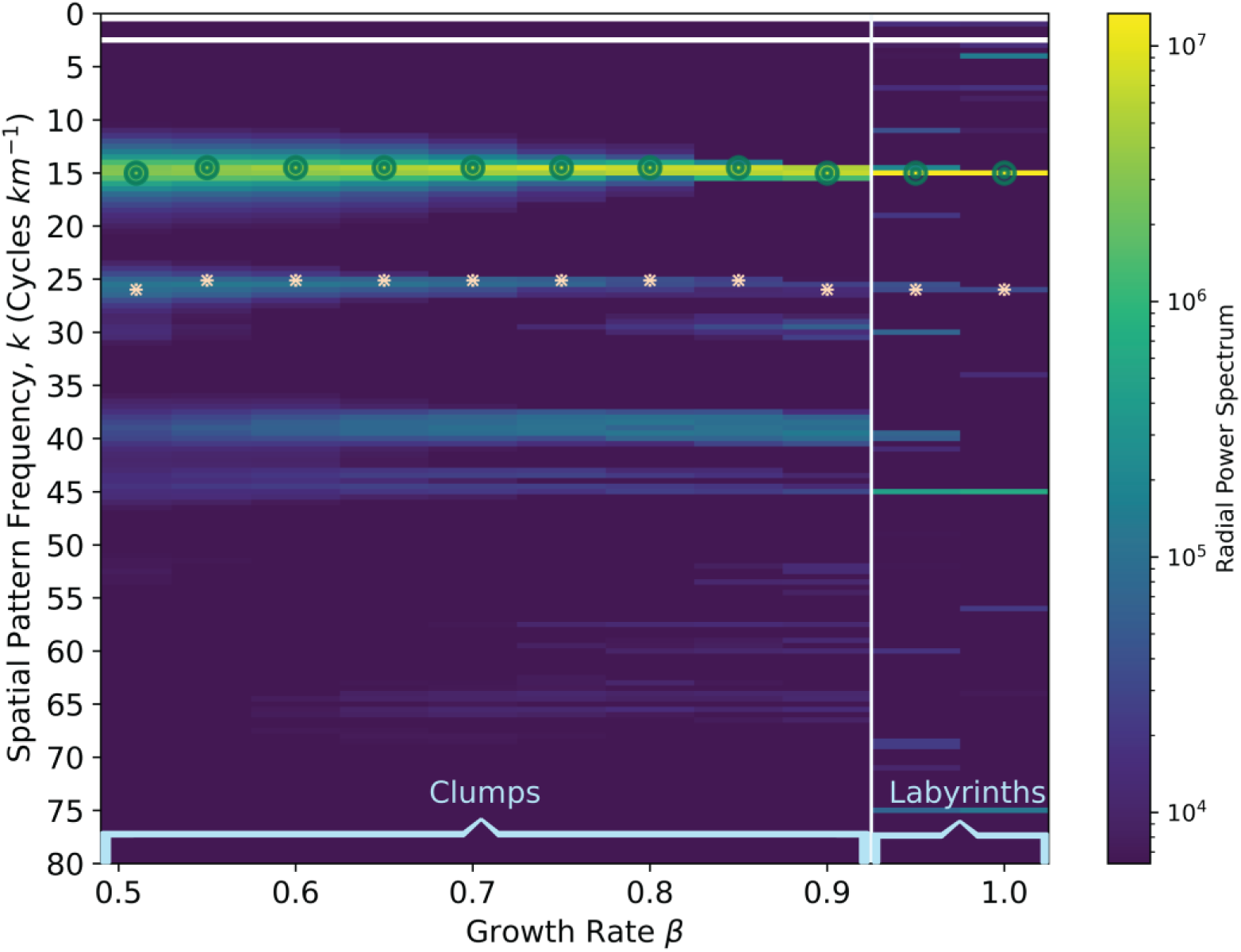
Heatmap for the radial power spectrum for the model in (Martinez-Garcia et al. 2013). Colors and axes like in Fig. 5 (main text). Note that the frequency at which the primary fundamental peak occurs does not change with growth conditions, differently from the Klausmeier model.

**Figure S6.3:**
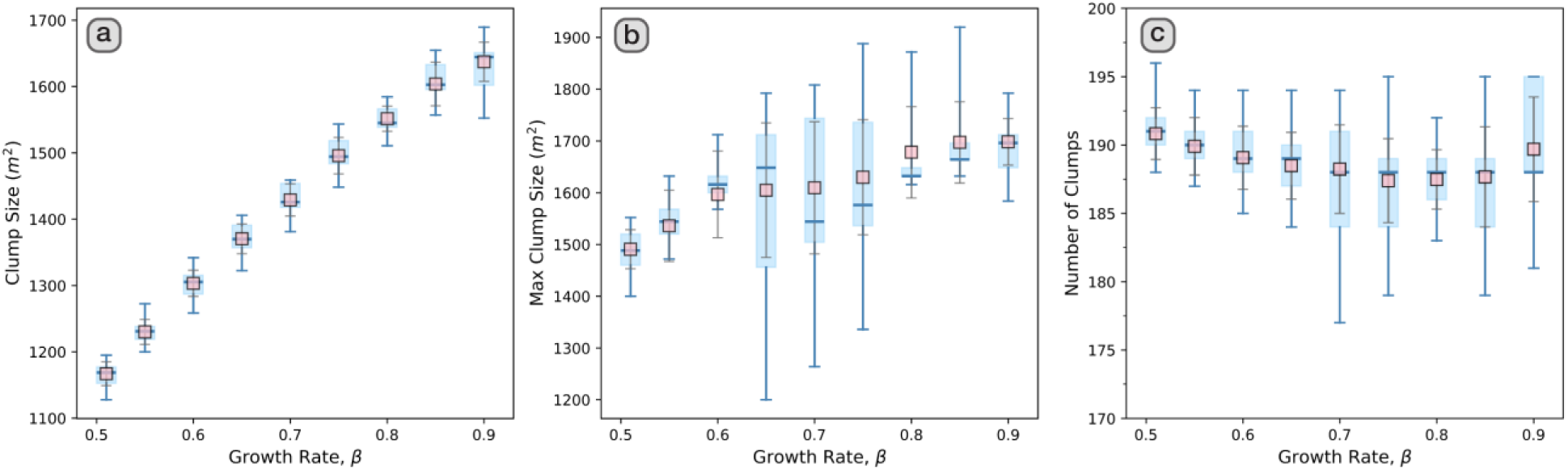
Average clump size (left), maximum clump size (center) and number of clumps (right) as a function of growth conditions for the model in (Martinez-Garcia et al. 2013).

#### ii) Model with spatial but no non-spatial bistability

The Rietkerk model (Rietkerk et al. 2002) is a three-equation model that, in addition to tracking vegetation density and surface water, also monitors the dynamics of soil water. Thus, its equations use terms that are more ecologically detailed than those in the Klausmeier model, although this increased realism comes with at the price expense of a much larger set of parameters.

Like in the Klausmeier model, in the Rietkerk model there is a range of rainfall values for which perturbing the desertic state triggers the emergence of clumps, evidencing the spatial bistability of vegetated and desert states within that range; on the other hand, when diffusivity is neglected and therefore space is not relevant (i.e. mean-field case), the homogeneous system does not show bistability (see Fig. S6.1).

We ran preliminary analyses to test whether these dynamical differences between the Klausmeier and the Rietkerk models affected the behavior reported in the main text, utilizing for the latter model parameter values that have been used in the past for arid regions in Africa (Bonachela et al. 2015; Rietkerk et al., 2002). As shown in Fig. S6.2, the behavior of the power spectrum as rainfall changes is qualitatively similar to that of the Klausmeier model (Fig. 5).

First, the spatial frequency at which the primary fundamental peak is located increases with rainfall. Moreover, the radial spectrum shows a secondary fundamental peak that differs from the harmonic of the primary fundamental peak, and the height of this peak and the harmonic change as rainfall increases. In addition, these changes correlate with a change in clump number (Fig. S6.3): as aridity increases (i.e. rainfall decreases), the number of clumps decreases. These similarities reinforce the idea of using the behavior of the primary and secondary fundamental peaks, and that of the number of clumps, as indicator for spatial bistability.

The average and maximum clump size, on the other hand, both show a marked decline, which departs from the negligible changes observed for the Klausmeier model and seems closer to the behavior observed for the model with a smooth transition. Note that this behavior is observed within the full spatial bistability regime which, for this model, extends up to rainfall levels *R* = 1.00 mm/d and includes all clump patterns. Outside the spatial bistability regime, clumps converge to labyrinths, with the corresponding increase in clump size and decrease in number. These results point to the presence of mean-field bistability as an important factor influencing the behavior of clump morphology, i.e. whether clump size is constant or increases with rainfall depends on the presence or absence of mean-field bistability.

**Figure S6.4:**
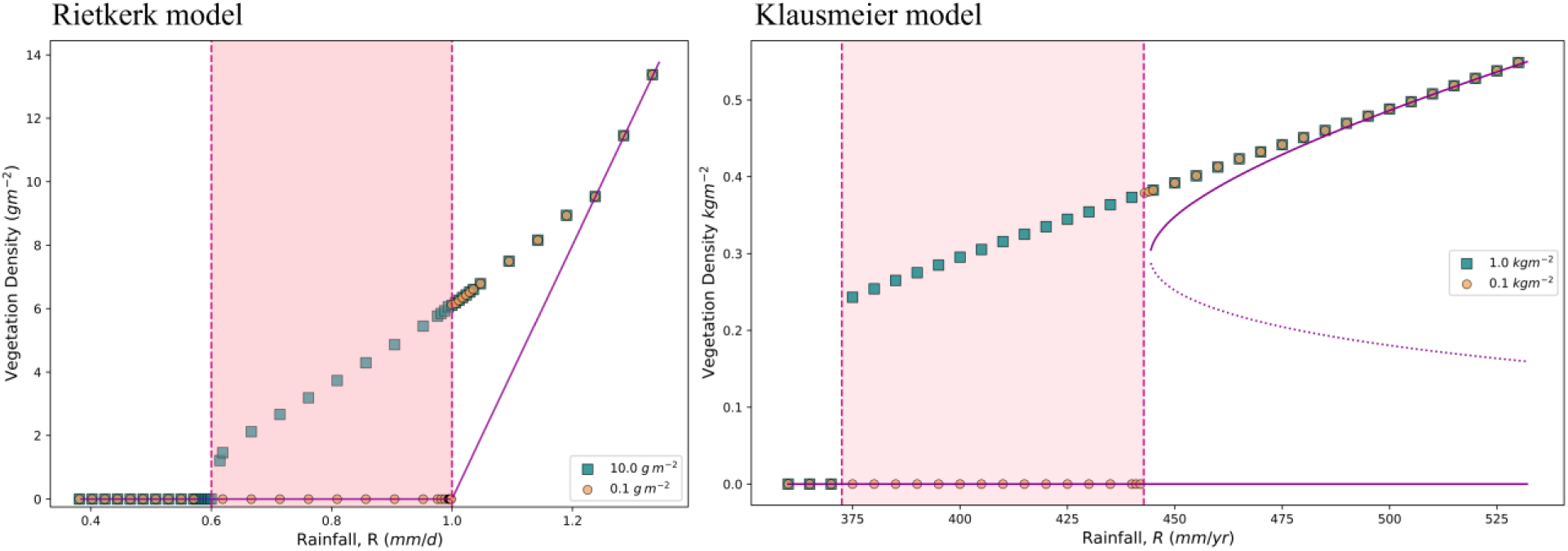
Phase diagram for the Rietkerk model (left) and, for reference, the Klausmeier model (right), for both the full model (points) and the model without diffusivity (i.e. non-spatial model, for which the result of the interactions between vegetation and water is a homogeneous state, purple lines). For the Rietkerk model (left), in the absence of patterns (i.e. when space is not relevant) the system does not show bistability but rather a smooth transition from vegetated to desertic state; note how the purple line only shows one value of vegetation for each rainfall value, as opposed to the two possible values shown by the purple line of the Klausmeier model (right). When considering vegetation and water dispersal (points), the Rietkerk model behaves like a scale-dependent feedback model and patterns develop after the homogeneous state is perturbed, giving rise to an abrupt desertification transition and spatial bistability for rainfall within [0.60,1.0] (shaded region). Differently from the Klausmeier model, in the Rietkerk model clumps are only observed within the spatial bistability region. For the non-spatial models (purple lines), the continuous transition in the Rietkerk model is also referred to as a transcritical bifurcation, and the discontinuous transition in the Klausmeier model as a fold bifurcation (where the dashed curve represents unstable values/states of the system).

**Figure S6.5:**
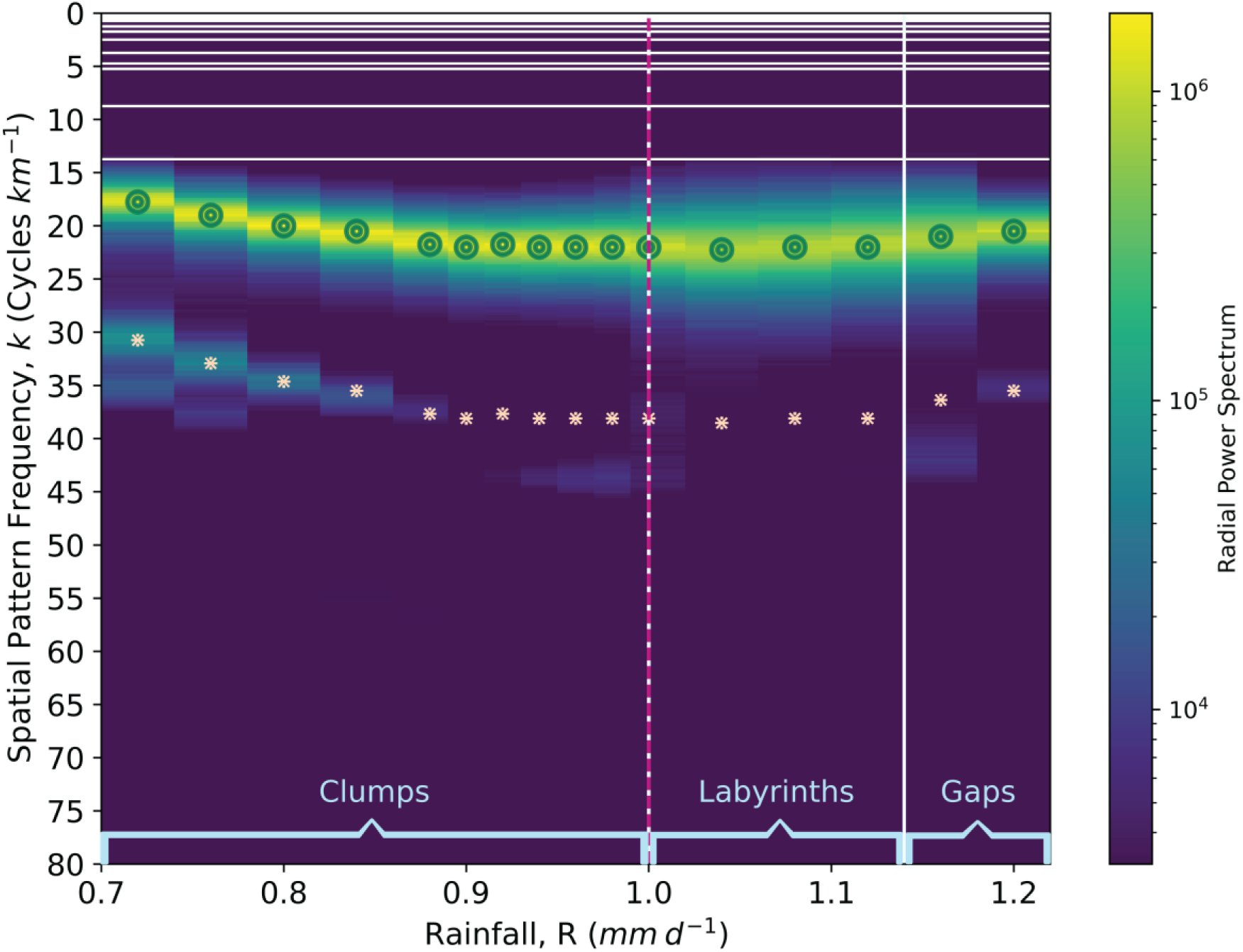
Heatmap for the radial power spectrum for the Rietkerk model. Colors and axes like in Fig. 5. Note the similarities with that figure for the Klausmeier model, with frequency at which the primary fundamental peak occurs changing with rainfall, and the existence of a secondary peak whose height decreases and ultimately becomes negligible as rainfall increases within the spatial bistability region.

**Figure S6.6:**
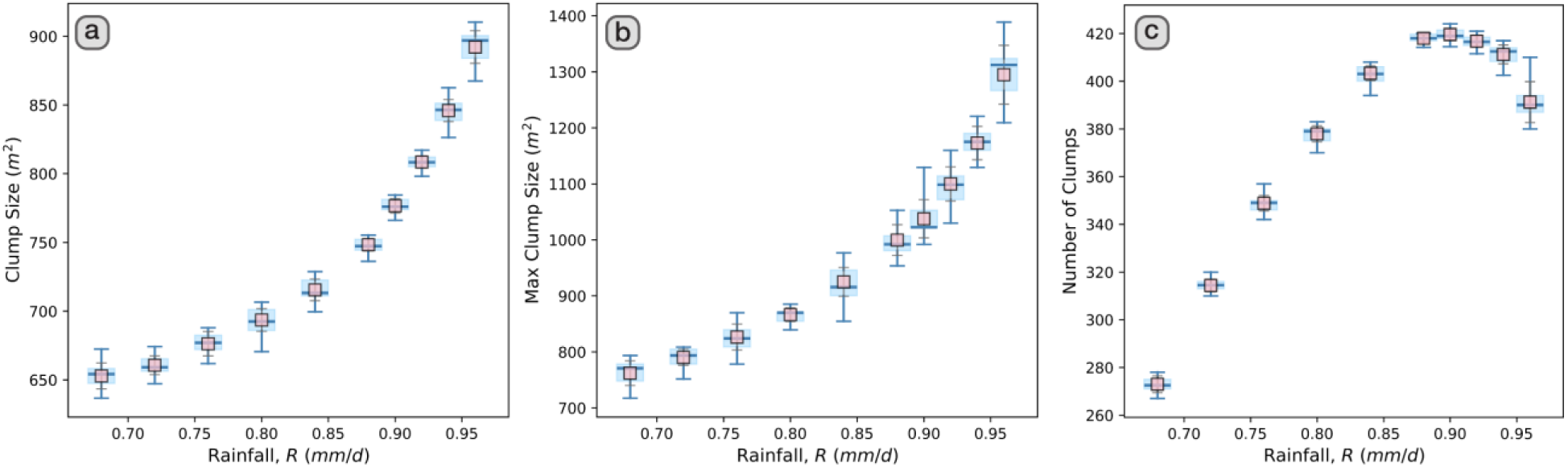
Average clump size (left), maximum clump size (center) and number of clumps (right) as a function of rainfall for the Rietkerk model. Note how, similarly to the Klausmeier model (and as expected from Fig. S6.5), the number of clumps decreases with lower rainfall; however, the decrease in clump size is marked, similarly to the behavior observed with the model in (Martinez-Garcia et al 2013).

#### iii) Model with both spatial and non-spatial bistability

To confirm the conclusions above, we simulated a version of the Rietkerk model in which surface water evaporation is also present (Siero, 2020). Like the Klausmeier model, this model has been shown to show both spatial and mean-field bistability. Unlike the Klausmeier model, however, both spatial and mean-field bistability are constrained to just a few values for rainfall levels. For this model, we used the same parametrization as the Rietkerk model, reducing by approximately a factor 3 all diffusivities and increasing by a factor 2 spatial resolution. We also used a high evaporation rate (0.1 d^-1^) that required a higher perturbation value. All together, we obtained the phase diagram in Fig. S6.7 showing both types of bistability, with a reduced spatial bistability regime that ends at a rainfall level (R=1.80 mm/yr) considerably lower than the point at which mean-field bistability starts (R∼2.1). Its corresponding heatmap behaves qualitatively like that of the Klausmeier model (Fig. S6.8). Importantly, although the number of clumps follows also the behavior of the Klausmeier model (and the classic Rietkerk model), clump size (both average and maximum) shows an initial constant behavior for low rainfall levels, but starts increasing around R∼1.80 (Fig. S6.9). Thus, this model confirms that the presence of mean-field bistability ultimately leads to constant clump size for high aridity (thus offering a potential early-warning signal for desertification), with clump size increasing with rainfall level as aridity goes away from the transition. This model also confirms that a constant number of clumps only results when the interactions in the system cannot produce spatial bistability.

**Figure S6.7:**
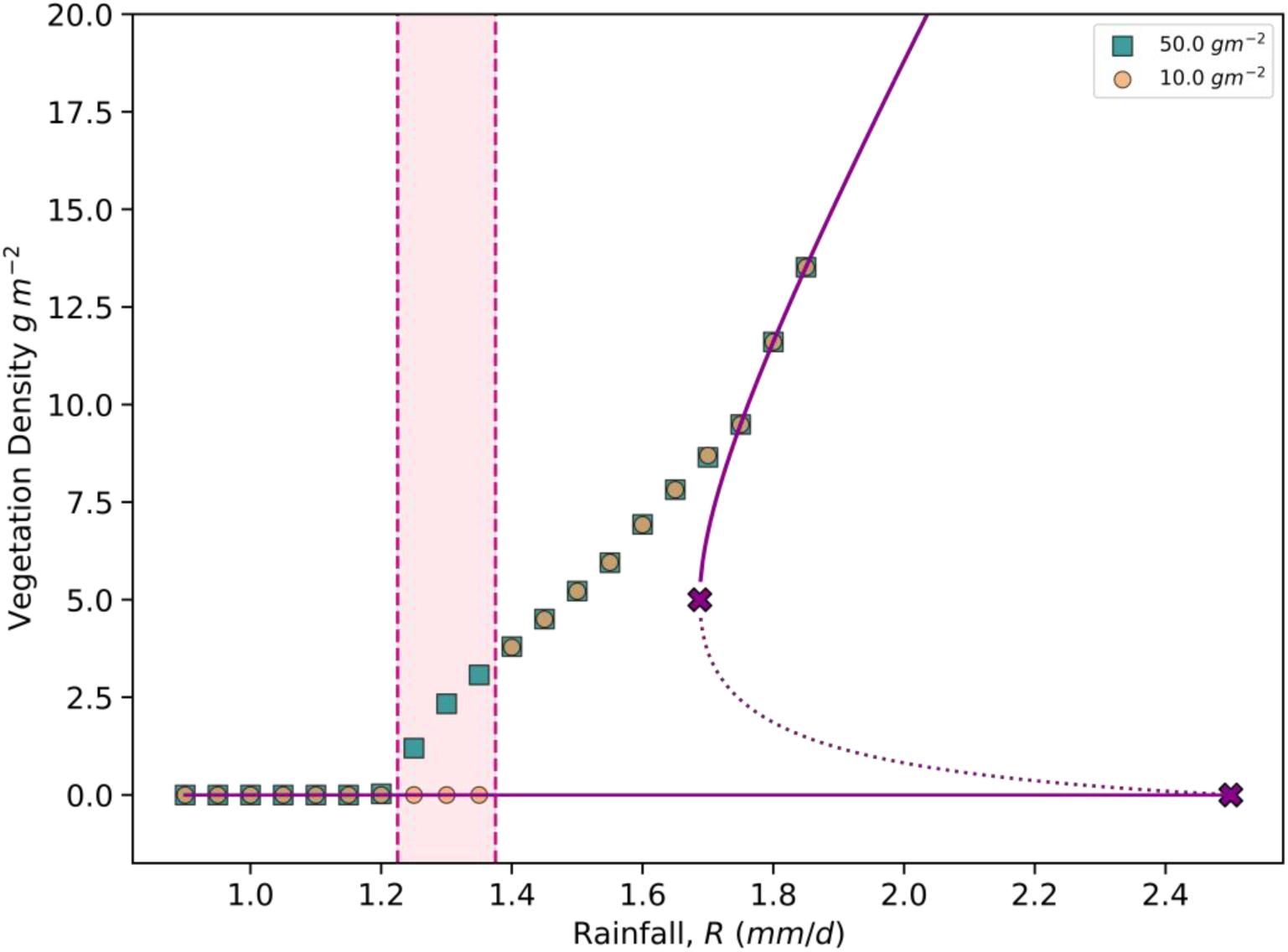
Phase diagram for the Rietkerk model with evaporation. As in the Klausmeier model (see e.g. Fig. S6.4, right panel), the system shows mean-field bistability (range of rainfalls between the two purple crosses). When considering vegetation and water dispersal, patterns develop after the homogeneous state is perturbed (perturbation values in legend), giving rise to an abrupt desertification transition and spatial bistability for rainfall within [1.25,1.35] (shaded region).

**Figure S6.8:**
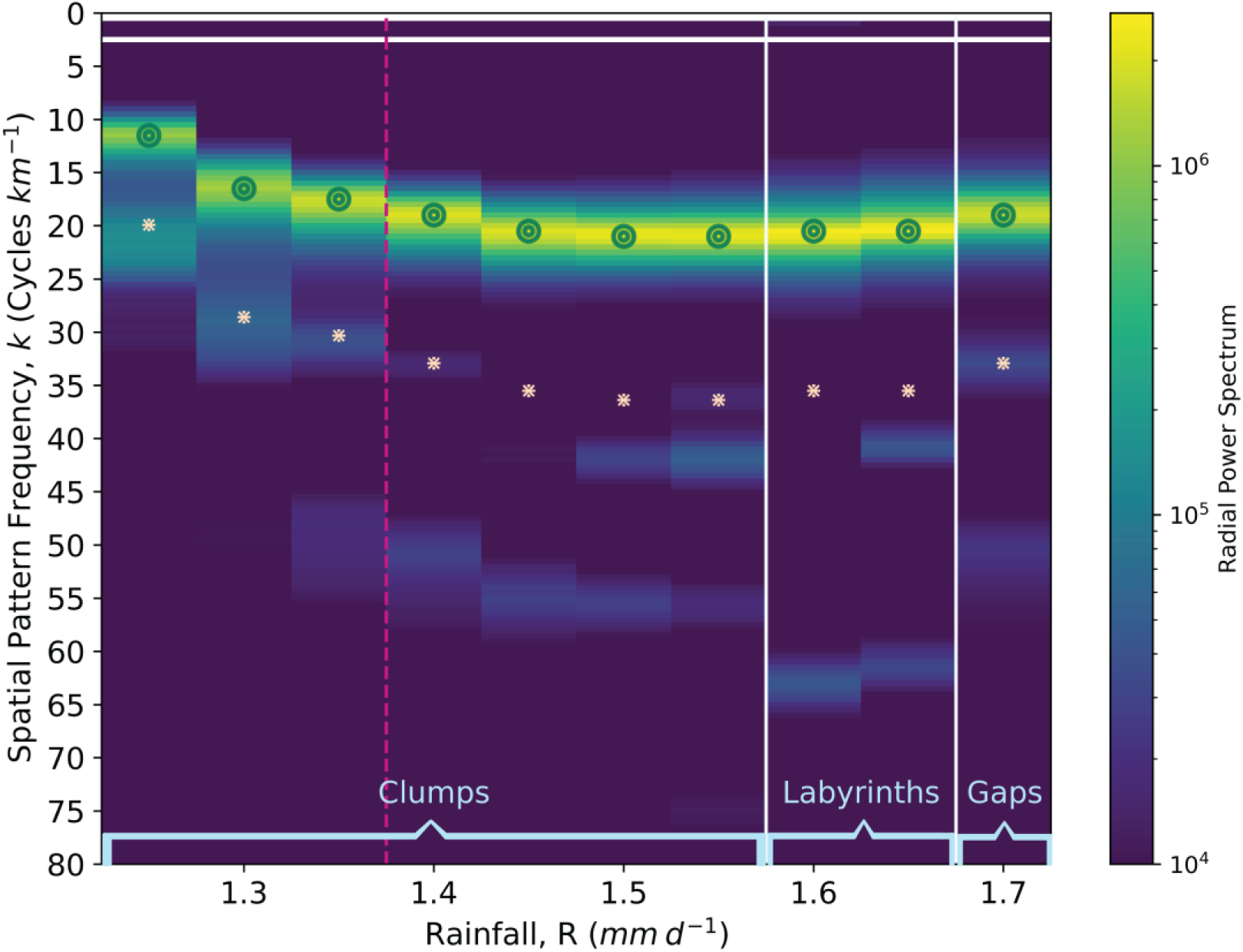
Heatmap for the radial power spectrum for the Rietkerk model with evaporation. Colors and axes like in Fig. 5. Note the similarities with that figure for the Klausmeier model, with frequency at which the primary fundamental peak occurs changing with rainfall, and the existence of a secondary peak whose height decreases and ultimately becomes negligible as rainfall increases within the spatial bistability region.

**Figure S6.9:**
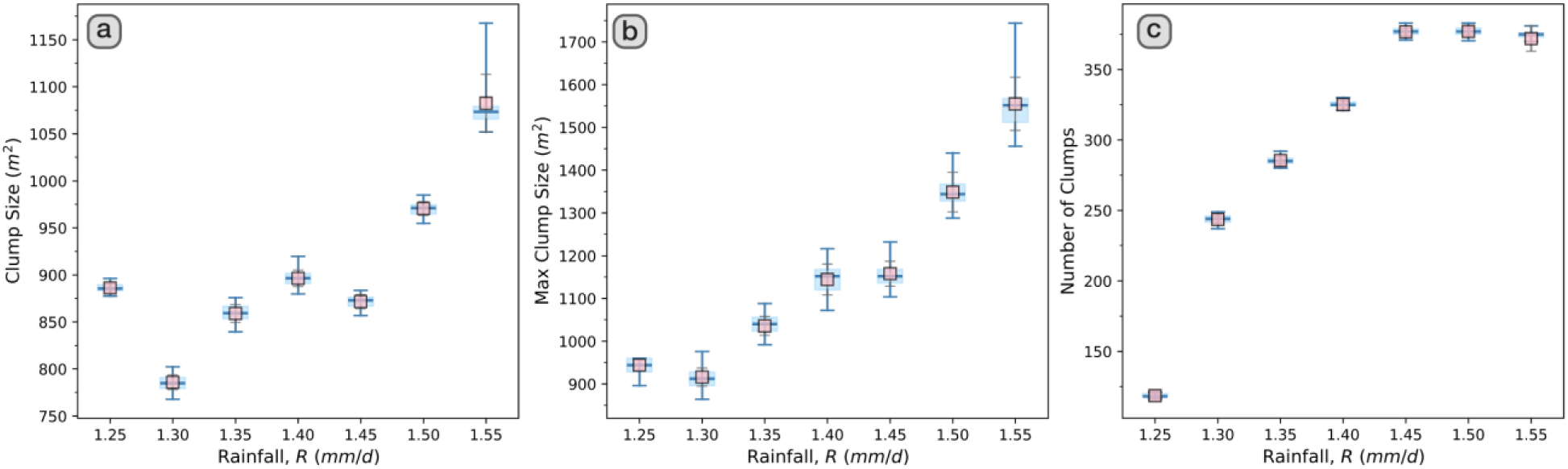
Average clump size (left), maximum clump size (center) and number of clumps (right) as a function of rainfall for the Rietkerk model with evaporation. Note how, similarly to the Klausmeier model, the number of clumps decreases with lower rainfall and the average and maximum size ultimately become constant.

## References

Bastiaansen R., Jaïbi, O., Deblauwe V, Eppinga M. B., Siteur K., Siero E., Mermoz S., Bouvet A., Doelman A., and Rietkerk M. (2018) Multistability of model and real dryland ecosystems through spatial self-organization, PNAS vol. 115, no. 44, 11256–11261.

Bastiaansen, R., Doelman, A., Eppinga, M. B and Rietkerk, M. (2020). The effect of climate change on the resilience of ecosystems with adaptive spatial pattern formation. Ecology Letters, 23(3), 414–429.

Borgogno, F., D’Odorico, P., Laio, F. and Ridolfi, L. (2009). Mathematical models of vegetation pattern formation in ecohydrology, Rev. Geophys.,47, RG1005.

Dakos, V., Kéfi, S., Rietkerk, M., van Nes, E. and Scheffer, M. (2011). Slowing down in spatially patterned ecosystems at the brink of collapse. The American Naturalist 177: E153–E166.

Deblauwe, V., Barbier, N., Couteron, P., Lejeune, O. and Bogaert, J. (2008). The global biogeography of semi-arid periodic vegetation patterns. Glob Ecol Biogoegr 17:715–723.

Deblauwe, V., Couteron, P., Lejeune, O., Bogaert, J. and Barbier, N. (2011). Environmental modulation of self-organized periodic vegetation patterns in Sudan. Ecography, 34, pp. 990–1001.

Eckhaus, W. (1965). Studies in non-linear stability theory. Springer.

Esri. (2009). World Imagery [Map]. Retrieved March 23, 2023, from https://www.arcgis.com/home/item.html?id=10df2279f9684e4a9f6a7f08febac2a9

Esri. (2013). Terrain: Slope in degrees [Map]. Retrieved March 23, 2023, from https://www.arcgis.com/home/item.html?id=af25a795273440deb449b336543602be

Gilad, E. (2004). Ecosystem engineers: From pattern formation to habitat creation, Phys. Rev. Lett., 93, 098105.

Gilad, E., von Hardenberg, J., Provenzale, A., Shachak, M. and Meron, E. (2007). A mathematical model of plants as Ecosystem Engineers. Journal of Theoretical Biology, 244(4), 680–691.

Hagos, M.G., (2001). The Influence of Tree Thinning and Subhabitat Differentiation on the Reproductive Dynamics of Acacia mellifera subsp. detinens. M.Sc. Agric-thesis, University of the Free State, Bloemfontein, South Africa.

Halder, K., & Bonachela, J. A. (2026a). Homogeneous States, Turing Instabilities, and Busse Balloons in Models for Spatial Self-Organization [Manuscript in preparation].

Halder, K., & Bonachela, J. A. (2026b). Spatial trophic dynamics shape and are shaped by desertification transitions [Manuscript under review].

Harris, C.R., Millman, K.J., van der Walt, S.J. et al. (2020). Array programming with NumPy. Nature 585, 357–362.

HilleRisLambers, R., Rietkerk, M., van den Bosch, F., Prins, HHT. and de Kroon, H. (2001). Vegetation pattern formation in semi-arid grazing systems. Ecology 82:50–61.

Intergovernmental Panel on Climate Change. (2022). Desertification. In Climate change and land: IPCC special report on climate change, desertification, land degradation, sustainable land management, food security, and greenhouse gas fluxes in terrestrial ecosystems (pp. 249–344). Cambridge University Press.

Kéfi, S., Rietkerk, M., Alados, C. et al. Spatial vegetation patterns and imminent desertification in Mediterranean arid ecosystems. Nature 449, 213–217 (2007).

Kéfi, S., Eppinga, M.B., de Ruiter, P.C. and Rietkerk, M. (2010). Bistability and regular spatial patterns in arid ecosystems. Theoretical Ecology, 3:257–269.

Klausmeier, C.A. (1999). Regular and irregular patterns in semiarid vegetation. Science 284, 1826–1828.

Leenaars, J. and Hengl, T. (2015) Africa SoilGrids - Texture. ISRIC.

Leenaars, J. and Hengl, T. (2017) Africa SoilGrids - Root zone plant available water holding capacity aggregated at ERZD. ISRIC.

Lejeune, O., Tlidi, M. and Couteron P. (2002). Localized vegetation patches: a self-organized response to resource scarcity. Phys. Rev. E 66: 010901R.

Lejeune, O., Tlidi, M. and Lefever, R. (2004). Vegetation spots and stripes: dissipative structures in arid landscapes. Int. J. Quantum Chem. 98:261–271.

Lenton, T. M., Armstrong McKay, D. I., Loriani, S., Abrams, J. F., Lade, S. J., Donges, J. F., Milkoreit, M., Powell, T., Smith, S. R., Zimm, C., Buxton, J. E., Bailey, E., Laybourn, L., Ghadiali, A., & Dyke, J. G. (2023). The global tipping points report 2023. University of Exeter. https://global-tipping-points.org/

Liu, F. T., Ting, K. M., & Zhou, Z.-H. (2008). Isolation forest. In 2008 Eighth IEEE International Conference on Data Mining (pp. 413–422). IEEE.

Manning, R., (1891). On the flow of water in open channels and pipes. Transactions of the Institution of Civil Engineers of Ireland. 20: 161–207.

Martínez-García, R., J. M. Calabrese, E. Hernández-García, and C. López (2013), Vegetation pattern formation in semiarid systems with-out facilitative mechanisms, Geophys. Res. Lett., 40, 6143–6147.

Martinez-Garcia, R., Cabal, C., Calabrese, J. M., Hernández-García, E., Tarnita, C. E., López, C. and Bonachela, J. A. (2023). Integrating theory and experiments to link local mechanisms and ecosystem-level consequences of vegetation patterns in drylands. Chaos, Solitons & Fractals, 166, 112881.

Meron, E. (2012). Pattern-formation approach to modelling spatially extended ecosystems. Ecol Model, 234, pp. 70–82.

Meron, E., Gilad, E., von Hardenberg, J., Shachak, M. and Zarmi, Y. (2004). Vegetation patterns along a rainfall gradient. Chaos Solitons Fractals 19: 367–376.

Namibia Agricultural Union, (2010). The Effect of Bush Encroachment on groundwater resources in Namibia, a Desk Top Study. Windhoek, Namibia: Namibia Agricultural Union. unpublished.

Okayasu, T. and Aizawa, Y. (2001). Systematic analysis of periodic vegetation patterns. Prog Theor Phys 106:705–720.

Pinto-Ramos, D., & Martínez-García, R. (2026). How spatial patterns can lead to less resilient ecosystems. Proceedings of the National Academy of Sciences, 123(14), e2511994123.

Rey, B., Fuller, A., Mitchell, D., Meyer, L. C. R., & Hetem, R. S. (2017). Drought-induced starvation of aardvarks in the Kalahari: An indirect effect of climate change. Biology Letters, 13, 20170301.

Rietkerk, M., Boerlijst, M.C., van Langevelde, F., HilleRisLambers, R., van de Koppel, J., Kumar, L., Prins, H.H.T. (2002). Self-organization of vegetation in arid ecosystems. Am Nat 160:524–530.

Rietkerk, M., Dekker, S.C., de Ruiter, P.C. and van de Koppel, J. (2004). Self-Organized Patchiness and Catastrophic Shift in Ecosystems. Science, 305, 1926–1929.

Rietkerk, M. and van de Koppel, J. (2008). Regular pattern formation in real ecosystems. Trends Ecol. Evol. 23, 169–175.

Rosenzweig, C., Elliott, J., Deryng, D., Ruane, A. C., Müller, C., Arneth, A., Boote, K. J., Folberth, C., Glotter, M., Khabarov, N., Neumann, K., Piontek, F., Pugh, T. A. M., Schmid, E., Stehfest, E., Yang, H., & Jones, J. W. (2014). Assessing agricultural risks of climate change in the 21st century in a global gridded crop model intercomparison. Proceedings of the National Academy of Sciences, 111(9), 3268–3273.

Scheffer, M., Bascompte, J., Brock, W.A., Brovkin, V., Carpenter, S.R., Dakos, V., … Sugihara, G. (2009). Early-warning signals for critical transitions. Nature, 461, pp. 53–59, 10.1038/nature08227

Sherratt, J.A., (2016). When does colonisation of a semi-arid hillslope generate vegetation patterns? J. Math. Biol. 73, 199–226.

Sherratt, J. A. and Lord, G. J. (2007). Nonlinear dynamics and pattern bifurcations in a model for vegetation stripes in semi-arid environments. Theor. Popul. Biol. 71: 1–11.

Siero, E. (2020). Resolving soil and surface water flux as drivers of pattern formation in Turing models of dryland vegetation: A unified approach. Physica D: Nonlinear Phenomena, 414, Article 132695.

Siteur, K., Siero, E., Eppinga, M.B., Rademacher, J.D., Doelman, A. and Rietkerk, M. (2014). Beyond Turing: The response of patterned ecosystems to environmental change. Ecol. Complex., 20, 81–96.

Snyder, K.A. and Tartowski, S.L. (2006). Multi-scale temporal variation in water availability: Implications for vegetation dynamics in arid and semi-arid ecosystems. Journal of Arid Environments, 65:219–234.

Thornthwaite, C. (1948). An Approach toward a Rational Classification of Climate. Geogr. Rev., 38(1), 55–94.

Ursino, N. (2005). The influence of soil properties on the formation of unstable vegetation patterns on hillsides of semiarid catchments, Adv. Water Resour., 28, 956–963.

Ursino, N. (2007). Modeling banded vegetation patterns in semiarid regions: Interdependence between biomass growth rate and relevant hydrological processes, Water Resour. Res., 43, W04412.

Vanderveken, L., Martínez Montero, M., and Crucifix, M. (2023). Existence and influence of mixed states in a model of vegetation patterns. Nonlinear Processes in Geophysics, 30(5), 585–599.

Veldhuis, M.P., Martinez-Garcia, R., Deblauwe, V. and Dakos, V. (2022). Remotely-sensed slowing down in spatially patterned dryland ecosystems. Ecography, e06139.

Von Hardenberg, J., Meron, E., Shachak, M., Zarmi, Y. (2001). Diversity of vegetation patterns and desertification. Phys. Rev. Lett. 87, 3–6.

Walther, G.-R. (2010). Community and ecosystem responses to recent climate change. Philosophical Transactions of the Royal Society B: Biological Sciences, 365, 2019–2024.

## References

Ahmed, K.R., Akter, S. (2017). Analysis of landcover change in Southwest Bengal Delta due to floods by NDVI, NDWI and K-means cluster with Landsat multi-spectral surface reflectance satellite data. Remote Sens Appl Soc Environ 8:168–181.

Alam, S. A., Starr, M. and Clark, B. J. F. (2013). Tree biomass and soil organic carbon densities across the Sudanese woodland savannah: A regional carbon sequestration study. Journal of Arid Environments, 89: 67–76.

Arcement, G. J. and Schneider, V. R. (1989). Guide for selecting Manning’s roughness coefficients for natural channels and flood plains. U.S. G.P.O.

Arthur, D. and Vassilvitskii, S. (2007). K-Means++: The advantages of careful seeding. Proc 18th Annu ACM-SIAM Symp Discrete Algorithms 1027–1035.

Bein, E., Habte, B., Jaber, A., Birnie, A., and Tengnas, B. (1996). Useful trees and shrubs in Eretria. Technical handbook No. 12, Regional Soil Conservation unit, Nairobi.

Bonachela, J.A., Pringle, R.M., Sheffer, E., Coverdale, T.C., Guyton, J.A., Caylor, K.K., Levin, S.A., Tarnita, C.E. (2015). Termite mounds can increase the robustness of dryland ecosystems to climatic change. Science 345:651.

Celik, T. (2009). Unsupervised change detection in satellite images using principal component analysis and K-means clustering. IEEE Geosci Remote Sens Lett 6:772–776.

Donaldson, C. H. (1969). Bush encroachment and methods of control with special reference to the Blackthorn (Acacia mellifera) problem of the Molopo area. Armoedsvlakte research station, Vryburg.

Esri. (2013a). World topographic map [Map]. Retrieved March 23, 2023, from https://www.arcgis.com/home/item.html?id=30e5fe3149c34df1ba922e6f5bbf808f

Esri. (2013b). Terrain: Slope in degrees [Map]. Retrieved March 23, 2023, from https://www.arcgis.com/home/item.html?id=af25a795273440deb449b336543602be

Halder, K., & Bonachela, J. A. (2026). Homogeneous states, Turing instabilities, and Busse balloons in dryland vegetation models [Manuscript in preparation].

Hoshen, J., Kopelman, R. (1976). Percolation and cluster distribution. I. Cluster multiple labeling technique and critical concentration algorithm. Phys Rev B 14:3438–3445.

Kittel, C., & McEuen, P. (2015). Introduction to solid state physics (8th ed.). John Wiley & Sons.

Sherratt, J. A. (2013). Numerical continuation of boundaries in parameter space between stable and unstable periodic travelling wave (wavetrain) solutions of partial differential equations. Advances in Computational Mathematics, 39, 175–192.

Leenaars, J. and Hengl, T. (2015) Africa SoilGrids - Texture. ISRIC.

Joubert, D.F., Rust, A., Smit, G., and Hoffman, M., (2017). Growth rates and mortality patterns of Acacia mellifera subsp. detinens in the semi-arid Highland Savanna, Namibia: Encroachment is not as rapid as previously believed. Namibian Journal of Environment, 1, A-5.

Joubert, D.F., Smit, G.N. and Hoffman, M.T. (2013). The influence of rainfall, competition, predation and germination and the establishment of an encroaching Acacia in an arid Namibian savanna. Journal of Arid Environments, 91: 7–13.

Maurer, C. R., Qi, R., & Raghavan, V. (2003). A linear time algorithm for computing exact Euclidean distance transforms of binary images in arbitrary dimensions. IEEE Transactions on Pattern Analysis and Machine Intelligence, 25(2), 265–270.

Olaode, A., Naghdy, G., Todd, C. (2014). Unsupervised classification of images: A review. Int J Image Process 8:2014–325.

Rousseeuw, P.J. (1987). Silhouettes: A graphical aid to the interpretation and validation of cluster analysis. J Comput Appl Math 20:53–65.

Seed, G. and Malik, A. (1968). Some aspects of measured and estimated evaporation in the Sudan, Durham theses, Durham University. Available at Durham E-Theses Online: http://etheses.dur.ac.uk/10121/

Turing, A. M. (1952). The chemical basis of morphogenesis. Philosophical Transactions of the Royal Society of London. Series B, Biological Sciences, 237(641), 37–72.

Ursino N (2005) The influence of soil properties on the formation of unstable vegetation patterns on hillsides of semiarid catchments. Adv Water Resour 28:956–963.

Walton, W. H. (1948). Feret’s statistical diameter as a measure of particle size. Nature, 162(4113), 329–330.

Wan, L., Li, Y., Cen, H., Zhu, J., Yin, W., Wu, W., Zhu, H., Sun, D., Zhou, W., He, Y. (2018). Combining UAV-based vegetation indices and image classification to estimate flower number in oilseed rape. Remote Sens 10:1484.

Zhang, T. Y., & Suen, C. Y. (1984). A fast parallel algorithm for thinning digital patterns. Communications of the ACM, 27(3), 236–239.

Zhao, J., Sang, Y., & Duan, F. (2019). The state of the art of two-dimensional digital image correlation computational method. Engineering Reports, 1(2), e12038.

